# Transcriptomic atlas of the morphologic development of the fungal pathogen *Coccidioides* reveals key phase-enriched transcripts

**DOI:** 10.1101/2024.10.13.618122

**Authors:** Christina M. Homer, Mark Voorhies, Keith Walcott, Elena Ochoa, Anita Sil

## Abstract

*Coccidioides spp*. are highly understudied but significant dimorphic fungal pathogens that can infect both immunocompetent and immunocompromised people. In the environment, they grow as multicellular filaments (hyphae) that produce vegetative spores called arthroconidia. Upon inhalation by mammals, arthroconidia undergo a process called spherulation. They enlarge and undergo numerous nuclear divisions to form a spherical structure, and then internally segment until the spherule is filled with multiple cells called endospores. Mature spherules rupture and release endospores, each of which can form another spherule, in a process thought to facilitate dissemination. Spherulation is unique to *Coccidioides* and its molecular determinants remain largely unknown. Here, we report the first high-density transcriptomic analyses of *Coccidioides* development, defining morphology-dependent transcripts and those whose expression is regulated by Ryp1, a major regulator required for spherulation and virulence. Of approximately 9000 predicted transcripts, we discovered 273 transcripts with consistent spherule-associated expression, 82 of which are *RYP1*-dependent, a set likely to be critical for *Coccidioides* virulence. ChIP-Seq revealed 2 distinct regulons of Ryp1, one shared between hyphae and spherules and the other unique to spherules. Spherulation regulation was elaborate, with the majority of 227 predicted transcription factors in *Coccidioides* displaying spherule-enriched expression. We identified provocative targets, including 20 transcripts whose expression is endospore-enriched and 14 putative secreted effectors whose expression is spherule-enriched, of which 6 are secreted proteases. To highlight the utility of these data, we selected a cluster of *RYP1*-dependent, arthroconidia-associated transcripts and found that they play a role in arthroconidia cell wall biology, demonstrating the power of this resource in illuminating *Coccidioides* biology and virulence.

## Introduction

*Coccidioides spp*. are dimorphic fungal pathogens found in the soil in the Southwest United States and other desert regions in Central and South America [1]. In the soil, they grow as hyphae that generate vegetative spores known as arthroconidia. Upon inhalation by a mammalian host, arthroconidia germinate and form a unique host-associated morphology known as the spherule [2]. Mature spherules rupture, releasing hundreds of internal cells known as endospores which can each go on to form another spherule in a cycle called spherulation. Notably, *Coccidioides* can cause infection in immunocompetent and immunocompromised individuals [3]. There is currently no cure for serious disseminated infections [4, 5]. Efforts to develop new treatments and prevention strategies have been hindered by a lack of molecular knowledge of the host form of *Coccidioides,* the spherule, including sparse sampling of the transcriptome during *Coccidioides* development. Prior studies have relied on microarray or low replicate number RNA-seq at one or two timepoints during spherule formation, in different conditions varying by laboratory, using different media to induce spherules versus hyphae, and only two published datasets profiled endospores after they have been released from spherules [6–12]. The spherule transcriptome remains under-characterized and the endospore transcriptome essentially unknown.

Despite limited molecular insight into spherulation, the Ryp1 transcription factor is known to be a major spherulation regulator [12]. Ryp1 is a WOPR-domain containing transcription factor whose orthologs (such as Wor1) in other fungi are regulators of morphology transitions and development [13–16]. Additionally, WOPR family proteins often regulate virulence factors [15–17] and are required for virulence in fungal pathogens [12, 18–21]. In *Coccidioides*, the *ryp1*Δ mutant is unable to form spherules and has an aberrant transcriptome in both spherule- and hyphal-conditions [12].

Here, we performed the first high-depth, high-density transcriptomic time courses of *Coccidioides* arthroconidia germinating into either hyphae or spherules that went on to release endospores. We leveraged the *ryp1*Δ mutant to define genes whose transcription is regulated by *RYP1* throughout these developmental trajectories, defined morphology-specific binding targets of Ryp1, and highlighted a particular role for *RYP1* in direct regulation of genes expressed in the spherule morphology. Additionally, we annotated transcription factors, identified candidate secreted effectors, and defined candidate endospore-associated genes. From these data, we selected a cluster of spore-associated genes that were *RYP1*-dependent and found that they play a role in arthroconidia cell wall biology, demonstrating the power of this transcriptomic atlas to uncover new biology. Together, these findings serve as a foundational resource for the study of this important fungal pathogen.

## Materials and Methods

### Strains and growth conditions

The wildtype *Coccidioides posadasii* strain Silveira (NR-48944) [22] was used for growth experiments and as the background for the generation of mutants. All manipulation of live *Coccidioides* strains were performed in a biosafety level 3 facility. Standard spherulation conditions: polypropylene flasks containing Converse media as previously published [23] were inoculated with 1 x 10^6^/mL arthroconidia (unless otherwise stated) and placed at 39°C, 10 % CO_2_, shaking at 120 rpm. Where noted, spherulation was induced under the same conditions with different media: DMEM (UCSF Media Core) containing 20 % FBS (Corning) or RPMI (UCSF Media Core) containing 10 % FBS. For hyphal growth, polypropylene flasks containing Converse media were inoculated with 1 x 10^6^ arthroconidia/mL and grown at 25**°**C (Figure 2, S2) or room temperature (RT) (Figure 4, S4) shaking at 120 rpm. All growth experiments were performed in 125 mL polypropylene flasks with 50 mL of media except for experiments in Figure 2, S2, which were performed in 1 L flasks with 350 mL of media (except for replicate 3 of *ryp1*Δ mutant in spherulation conditions, which was placed in 300 mL of media given limited arthroconidia stock, to maintain the same concentration across all samples) and Figure 4, S4, which were performed in 1 L flasks with 330 mL Converse for wildtype and 500 mL flasks with 100 mL Converse for the *ryp1*Δ mutant.

### Microscopy of spherules and hyphae

At stated timepoints for light microscopy, cells were fixed in 4 % paraformaldehyde (Electron Microscopy Sciences) at RT for 30 minutes and washed twice in PBS, pelleting cells by centrifugation for 2 minutes at maximum speed between washes. Cells were visualized using 40X DICII objective on a Zeiss Axiovert 200 microscope, with additional 1.6x Optovar magnification.

### *ryp1*Δ deletion mutant generation

The *ryp1*Δ deletion mutant was created as previously described [24]. In brief, Phusion polymerase (Fisher) was used to amplify the hygromycin selection cassette (sequence from pMAD91 [25]) using Primer 1 and Primer 2 (Table S1), with 50 bp sequence complementary to the 5’ and 3’ flanking regions of the *RYP1* gene, D8B26_000722. These primers were used to generate the initial template and, given low efficiency, another round of amplification was performed with Primer 3 and Primer 4 at Tm 58.5°C. 2 µg of repair template DNA was gel extracted and purified using the Qiagen gel extraction kit and concentrated by isopropanol precipitation for transformation. Protoplasts were generated as previously described [26] with minor alterations: 100 mL of liquid 2x GYE media (2 % Dextrose (Fisher), 1 % Bacto Yeast Extract (Gibco)) were inoculated with 5 x 10^8^ arthroconidia and incubated shaking at 140 rpm, 30°C for ∼18h until germ tubes were visible by light microscopy. Cells were then centrifuged at 2800 x g for 10 minutes at RT, washed twice in 15 mL osmotic buffer A (OBA: 50 mM potassium citrate (Sigma), 0.6 M KCl (Fisher) at pH 5.8) and resuspended in cell wall digestion buffer (*Trichoderma harzanium* lysing enzymes 4 mg/ml (Sigma), Driselase from *Basidiomycetes* 7.5 mg/ml (Sigma) in OBA). Cell wall digestion was performed by shaking platform at 50 rpm, 30°C for 70 minutes. Protoplasts were pelleted by centrifugation at 900 x g for 10 minutes at RT and resuspended in osmotic buffer B (OBB at pH 5.8: 10 mM sodium phosphate (Fisher), 1.2 M MgSO_4_ (Fisher)). Trapping buffer at pH 7.5 (100 mM MOPS (Sigma), 0.6 M sorbitol (Sigma)) was overlaid on top of OBB and phase separation established through 15 minute centrifugation at 2800 x g at RT. Protoplasts were recovered from the interphase at RT and diluted 1:10 into MOPS buffer containing sorbitol at pH 6.5 (10 mM MOPS (Sigma), 1 M sorbitol (Sigma)). Protoplasts were pelleted by centrifugation at 900 x g for 10 minutes at RT and washed twice in MOPS buffer containing sorbitol and calcium (MSC buffer at pH 6.5: 10 mM MOPS (Sigma), 1 M sorbitol (Sigma), 20 mM CaCl_2_ (Fisher)). Cas9 ribonucleoprotein complexes (purchased from IDT) targeting each end of *RYP1* were assembled in vitro immediately before use as previously described [27] using the crRNA sequences 1 and 2 (Table S1) and the universal Alt-R tracrRNA (IDT). Ribonucleoprotein complexes, 2 µg of repair template DNA, and ∼10^7^ protoplasts in 100 µL MSC buffer were mixed with 30 µL 60 % PEG 3350 (Spectrum) and incubated on ice for 30 minutes. 900 µL of 60 % PEG was added followed by an additional 30 minutes of incubation at RT. Protoplasts were pelleted at 8000 rpm for 15 minutes at RT, followed by discarding 500 µL of supernatant, then an additional 2 minutes of centrifugation at 8000 rpm at RT, and removal of the remaining supernatant. The protoplast pellet was resuspended in 500 µL of MSC buffer and combined with melted GYES soft agar (1 % dextrose, 0.5 % Bacto yeast extract, 1 M sucrose (Sigma), 0.7 % Bacto-Agar (BD)) cooled to 46°C and overlaid onto a pre-warmed GYES agar plate (1 % dextrose, 0.5 % Bacto yeast extract, 1 M sucrose (Sigma), 2 % Bacto-Agar). Plates were incubated at 30°C for 48h. GYE soft agar (1 % dextrose, 0.5 % Bacto yeast extract, 0.7 % Bacto-Agar) with 75 µg/mL hygromycin (Invitrogen) was overlaid on colonies and plates were incubated for an additional 5-7 days at 30°C until colonies appeared on the surface of the agar. Single colonies were transferred to 2x GYE plates with 75 µg/mL hygromycin and grown again at 30°C. Colonies were passaged to fresh 2x GYE plates with 75 µg/mL hygromycin every 5-7 days for 9 generations.

### *DitCluster*Δ and *DitClusterSmall*Δ deletion mutant generation

*DitCluster*Δ mutants were generated through the same procedure as described above except for the following alterations: 333 ng of synthesized repair template (Table S1, Azenta) was used instead of generating this by PCR. Protoplasting was done similarly but omitting the germ tube washes with OBA, digested with enzyme mixture for 40 minutes total, and omitting the 2 final washes in MSC buffer. Ribonucleoprotein complexes were assembled as described above using crRNA sequences 3 and 4 for *DitCluster*Δ and crRNA sequences 3 and 5 for *DitClusterSmall*Δ. 6 x 10^5^ protoplasts in 100 µL MSC buffer were mixed with 25 µL 60 % PEG and incubated on ice for 50 minutes. The remainder of the transformation was done as described above. Colonies were passaged on 2x GYE plates with hygromycin for a total of 4 generations.

### gDNA Extraction and *ryp1Δ/DitCluster*Δ/*DitClusterSmall*Δ mutant verification

Hyphae were scraped from a colony and submerged in 700 µL lysis buffer (50 mM Tris pH 7.2 (Fisher), 50 mM EDTA (Fisher), 3 % SDS (Fisher), 1 % 2-Mercaptoethanol (Biorad)) and bead beat for 2 minutes at maximum speed (Biospec Mini Beadbeater), then incubated for 1 hour at 65°C after which 800 µL phenol/chloroform/isoamyl alcohol (Thermo) was added to each tube and mixed by inverting several times. Tubes were centrifuged at maximum speed for 15 minutes and genomic DNA was precipitated from the aqueous phase with 2-propanol (Thermo) and 0.13 M sodium acetate (Thermo), then pelleted by centrifugation at maximum speed for 2 minutes. The DNA pellets were washed twice with 70 % ethanol (Fisher) then dried at 50°C for 5-10 minutes. DNA was eluted into TE+RNAse A (0.15 mg/mL, Qiagen) then stored at −20°C until use. Mutant verification was accomplished by PCRs with primers (Table S1) designed to query correct integration of the repair cassette at the 5’ (Primers 5/6 for *ryp1*Δ and Primers 11/12 for *DitCluster*Δ and *DitClusterSmall*Δ) and 3’ (Primers 7/8 for *ryp1*Δ, Primers 13/14 for *DitCluster*Δ, Primers 14/15 for *DitClusterSmallΔ)* ends of the cassette, and loss of the native gene sequence (Primers 9/10 for *ryp1*Δ, Primers 16/17 for *DitCluster*Δ and *DitClusterSmall*Δ, and additionally Primers 17/18 for *DitCluster**Δ***). Mutants were further verified by whole genome sequencing through SeqCenter (Illumina Whole Genome Sequencing, 2GB coverage). Reads were aligned to the reference with BWA MEM 0.7.17 [28] using default settings, bedgraph files were generated using BEDTools 2.30.0 [29] and Integrative Genome Viewer [30] was used to visualize resulting coverage. This procedure verified loss of the coding sequence for each intended gene deletion and insertion of the hygromycin cassette at the site of the deleted gene with no off-target hygromycin insertions (by analyzing the position of discordant reads where one read in a pair mapped to the hygromycin cassette). Since we had one isolate of the *ryp1*Δ mutant, we further audited that mutant as follows: PILON 1.23 [31] was used to generate a reference guided assembly of the *ryp1*Δ mutant genome from paired-end Illumina reads and the GCA_018416015.2 *Coccidioides posadasii* reference [22]. BWA was used to generate alignments as specified above and those alignments were used as input to PILON in variant mode. This procedure was repeated for 15 iterations and able to fully assemble the hygromycin cassette insertion that replaced *RYP1* (*ryp1*Δ mutant genome fasta file in Supplementary Materials).

### Arthroconidia generation

Arthroconidia from wildtype or the *ryp1*Δ mutant were inoculated onto 2x GYE agar with Penicillin/Streptomycin (100 U/mL penicillin and 100 g/mL streptomycin, UCSF Media Core) or 2x GYE agar with 75 µg/mL hygromycin, respectively, in T225 tissue culture flasks and grown for 4-6 weeks at 30°C, until the hyphal mat appeared dry and flattened as previously described [32]. Arthroconidia were harvested 0-2 days prior to initiating spherulation and stored at 4°C until use. Arthroconidia harvest was done as previously described [32], by adding PBS (UCSF Media Core) to tissue culture flasks with the hyphal mat, scraping to resuspend and filtering through a 70-micron mesh filter. Arthroconidia were then washed twice with PBS and resuspended in PBS at appropriate concentrations for downstream assays. Arthroconidia were quantified using a plastic hemacytometer sealed with nail polish.

### RNA extraction and RNA-Seq library preparation

RNA from arthroconidia were collected from the same arthroconidia stock in triplicate by placing 5 x 10^7^ arthroconidia into Trizol LS (Ambion) and bead beating for 2 minutes. For all other samples, at indicated timepoints, RNA was extracted by pelleting cells by centrifugation at 1200 x g for 5 minutes at RT, removing supernatants, and flash freezing cell pellets in liquid nitrogen. Cell pellets were resuspended in Trizol, thawed, and bead beat for 2 minutes. Samples were stored at −80°C until all samples at all timepoints in an individual experiment had been collected. RNA was extracted using the Direct-zol RNA Miniprep Plus isolation kit (Zymo) with on-column DNAse digestion step extended for 15-30 minutes. Sequencing libraries were prepared using the NEBNext polyA mRNA magnetic isolation module and NEBNext Ultra II Directional RNA Library Prep kit with dual-indexed multiplexing barcodes. Library quality and adapter dimer contamination was analyzed using Agilent Bioanalyzer High Sensitivity DNA Chips. An additional round of library size selection was performed using homemade Serapure size selection beads [33] for libraries containing significant adapter dimers. Final library concentrations were measured using the Qubit High Sensitivity or Broad Range reagents depending on estimated library concentration by Bioanalyzer analysis of libraries. Libraries were pooled and sequencing was performed on a single lane of the HiSeq 4000 at the Center for Advanced Technology (UCSF) (Figure 1), or on 2 lanes of Novaseq S2 (Figure 2)/NextSeq 2000 P3 (Figure S4) at the Chan Zuckerberg Biohub-San Francisco.

**Figure 1:**
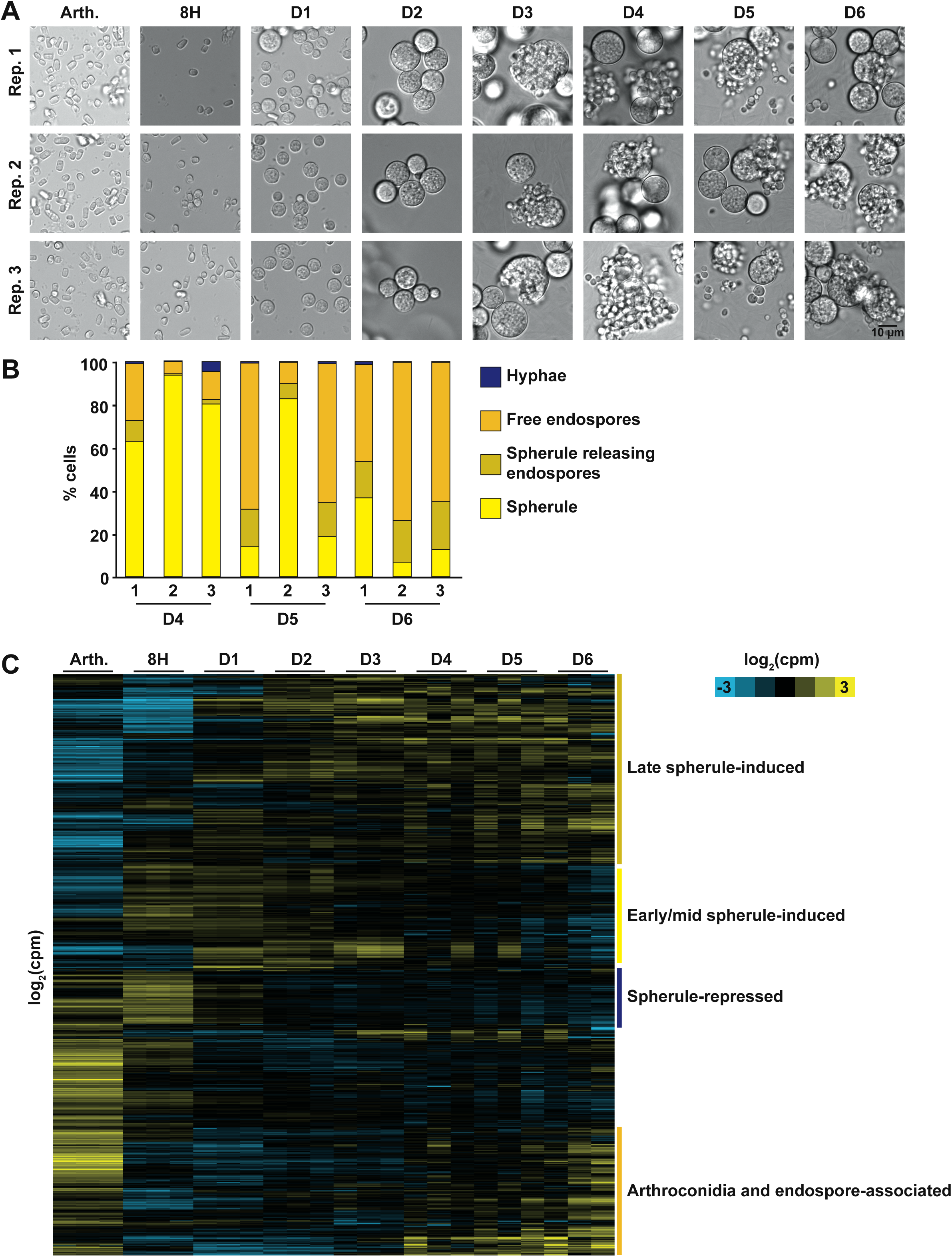
The transcriptome of arthroconidia germinating into spherules and releasing endospores. A. Micrographs of fixed samples from each flask at the time of RNA harvest. Endospore release was first observed on day 3. B. Quantification of the proportion of each morphology in cultures on days 4-6. n ≥ 10 fields of view counted for each sample. C. Heatmap of transcript abundance as arthroconidia germinate into spherules and release endospores. Transcripts that changed at least 2-fold between any adjacent timepoint or for a timepoint compared to arthroconidia (with 5 % FDR per limma) are included as mean centered rows in this heatmap. Rows are clustered based on correlation across all columns. Log_2_(counts per million) indicated by yellow and blue shading.

**Figure 2:**
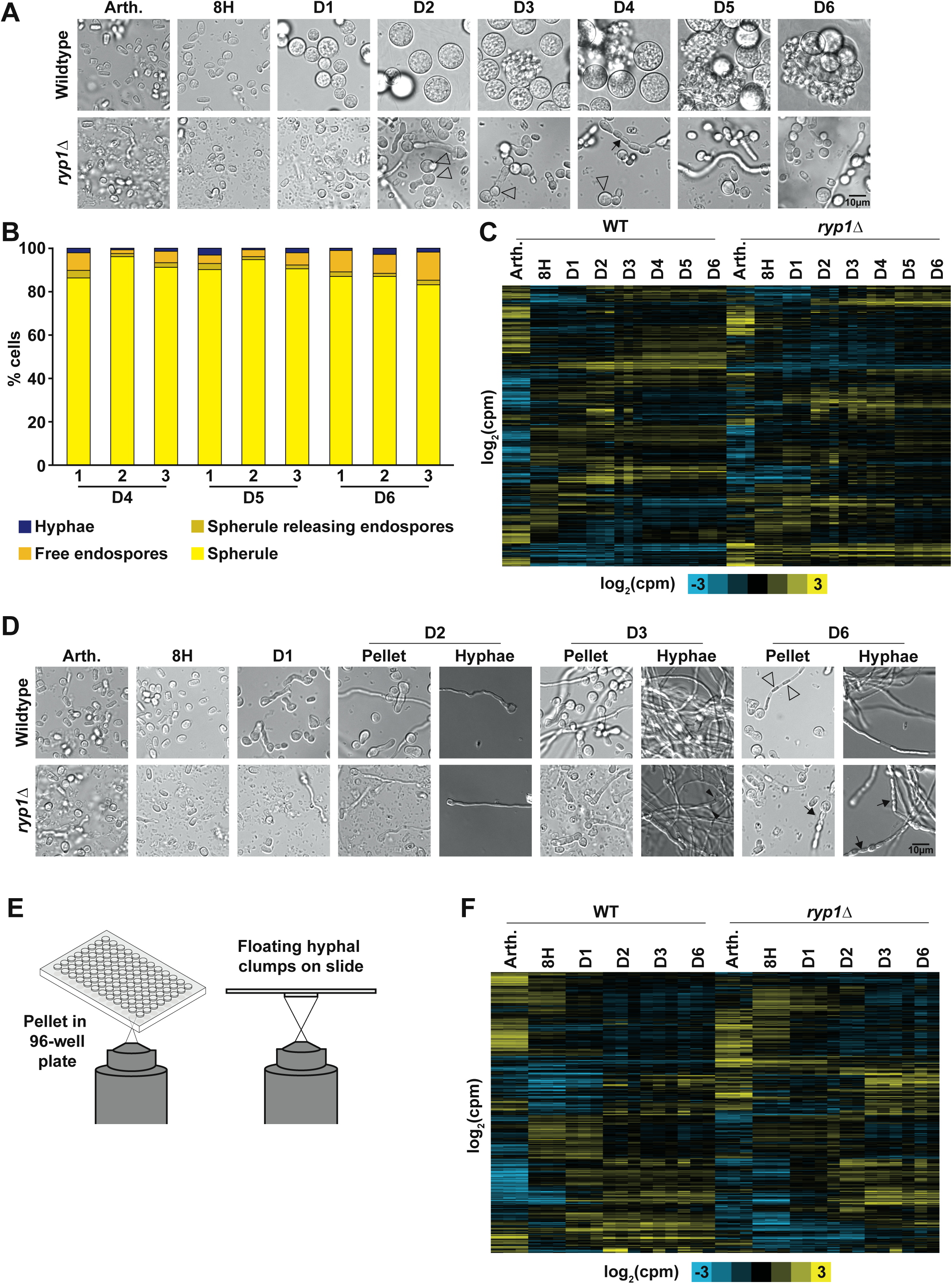
Spherule and hyphal transcriptomes are dependent on *RYP1*. A. Micrographs of fixed samples from each flask at the time of RNA harvest for one replicate of spherule growth in Converse, 39°C, 10 % CO_2_. Subsequent samples were taken from the same flask over time. Endospore release was first observed on day 3 in wildtype. As expected, the *ryp1*Δ mutant did not form spherules under these conditions. Surprisingly, it did form smaller rounded structures of unclear significance (open arrowheads) in addition to hyphae and some chains of oblong cells (black arrows) which have not been reported previously in *Coccidioides* to our knowledge. B. Quantification of the proportion of each morphology in cultures on days 4-6, n ≥ 20 fields of view counted for each sample. C. Heatmap of transcript abundance over time in spherulation conditions. Transcripts that were significantly differential (5 % FDR, 2-fold difference between any adjacent timepoints, a timepoint compared to arthroconidia, or between wildtype and the *ryp1*Δ mutant at any paired timepoint) are included as mean centered rows in this heatmap. Rows are clustered based on correlation across all columns. Log_2_(counts per million) indicated by yellow and blue shading. D. Micrographs of fixed samples from each flask at the time of RNA harvest for one replicate of hyphal growth in Converse, 25°C. Subsequent samples taken from the same flask over time. ‘Pellet’ and ‘Hyphae’ are the same biological samples prepared in different ways as described in E. Open arrows indicate hyphae forming initial arthroconidia. Black arrowheads indicate branching hyphae. Black arrows indicate chains of oblong cells similar to those observed for *ryp1*Δ in spherulation conditions. E. Schematic of preparation of hyphal samples for microscopy. ‘Pellet’ samples were placed in a 96-well plate with glass bottom and pelleted at 584 x g for 2 minutes prior to visualization. For ‘hyphal’ samples, 5µL of fixed samples containing small clumps of hyphae were placed on a slide with a coverslip prior to visualization. F. Heatmap of transcript abundance over time in hyphal conditions, displayed in the same manner and with the same criteria for inclusion as in C.

### RNA-seq data analysis

Analysis was conducted as previously described [12] with alterations below. Briefly, estimated counts of each transcript were calculated for each sample by alignment-free comparison against the predicted mRNA for the published Silveira genome [22] using KALLISTO version 0.46.2 [34]. Further analysis was restricted to transcripts with raw counts ≥ 10 in at least one sample across an individual experiment. Differentially expressed genes were identified by comparing replicate means for contrasts of interest using LIMMA version 3.30.8 [35]. Genes were considered significantly differentially expressed if they were statistically significant (at 5 % FDR) with an absolute log_2_ fold change ≥ 1 for a given contrast unless otherwise noted in the text.

### Ryp1 ChIP-Seq

50 mL of cultures were collected at the start of the experiment (arthroconidia), 8 hour, day 1, day 2, and day 4 timepoints from spherule or hyphal growth induced as described above. Paired samples for RNA-seq (50 mL initial culture for arthroconidia and 8 hour spherules/hyphae timepoints, then 10 mL of day 1, 2, and 4 spherules/hyphae timepoints) were also collected and processed as above. Cells were immediately crosslinked with 1 % formaldehyde (Neta Scientific) and incubated at RT for 20 minutes, mixing every 4 minutes. Crosslinking was then quenched with 125 mM glycine (Fisher), and samples were incubated for 5 minutes at RT, then frozen at −80°C. Frozen samples were collected for all timepoints in the experiment prior to downstream processing. All of the following buffers were made using autoclaved ddH_2_O in baked glassware/DNA-free plastic tubes. Samples were thawed, pelleted, and washed twice with 25 mL of TBS (20 mM Tris-HCl pH 7.5 (Fisher), 150 mM NaCl (EMD)), centrifuging at 3000 x g for 5 minutes at 4°C at each step. Pellets were resuspended in 700 µL of 4°C lysis buffer (50 mM HEPES (Fisher) / KOH (Fisher), 140 mM NaCl, 1 mM EDTA (Fisher), 1 % Triton X-100 (Acros Organics), 0.1 % sodium deoxycholate (Sigma), 2X Halt Protease Inhibitor Cocktail (ThermoFisher), 0.2X Halt Phosphatase Inhibitor Cocktail (ThermoFisher)). Cells were lysed by 8 x 1 minute cycles of bead beating (0.5 mm Zirconia/silica beads (Biospec)) at RT with 2 minute rests on ice in between each cycle. Insoluble chromatin was pelleted by centrifugation at 8000 rpm for 10 minutes at 4°C, resuspended in 350 µL cold lysis buffer, and then sonicated (Diagenode Biorupter) for 15 cycles (30 seconds on, 30 seconds off). Cell debris was removed by centrifugation at 14000 rpm at 4°C, yielding the soluble chromatin fraction. 10 µL of input DNA was removed from the sample and placed in TE (10 mM Tris HCl pH 8.0, 1 mM EDTA) with 1 % SDS (Fisher). The remaining chromatin was immunoprecipitated (IP) with 5 µg of a polyclonal rabbit antibody against an epitope of Ryp1 (ID: 3878, Epitope: VYRELDKPFPPGEKKRAMKK, Bethyl laboratories) with rotation overnight at 4°C. 50 µL of a 50 % slurry of protein A dynabeads (Life Technologies, washed 2x with cold TBS and 3x with cold lysis buffer) were added to each protein/antibody mixture and incubated an additional 3 hours at 4°C with rotation. Beads were then pelleted on a magnetic rack and washed 2x with cold lysis buffer (without protease or phosphatase inhibitors), 2x with cold lysis buffer with 500 mM NaCl instead of 140 mM NaCl, 2x with cold wash buffer (10 mM Tris-HCl pH 8.0, 250 mM LiCl (Sigma), 0.5 % NP-40 (Fluka), 0.5 % sodium deoxycholate, 1 mM EDTA), and 1x with cold TE. Bound protein/DNA complexes were eluted by adding 110 µL of elution buffer (50 mM Tris-HCl pH 8.0, 10 mM EDTA, 1 % SDS), vortexing, and incubating 10 minutes at 65°C, mixing every 2 minutes. Samples were placed on a magnet and 100 µL eluate was moved to a new tube. Then, 150 µL TE with 0.65 % SDS was added to the same beads, vortexed and placed back on the magnet, allowing 150 µL to be removed and combined with previous eluate for 250 µL for each sample. 1 µL of proteinase K (20 mg/mL, Qiagen) was added to these IP samples as well as the previously collected input samples. All samples were incubated at 65°C overnight (approximately 16 hours). 2 µg of RNase A (Qiagen) were added to each sample, which was incubated at 37°C for 1 hour. Samples were then purified using the Zymo ChIP DNA Clean and Concentrator kit. Libraries were prepared using the NEBNext Ultra II DNA Library Prep kit, with an additional round of higher and lower size selection (calibrated to select for 150-350 bp fragment sizes) using homemade Serapure size selection beads. Library quality and adapter dimer contamination was analyzed using Agilent Bioanalyzer High Sensitivity DNA Chips. Final library concentrations were measured using the Qubit High Sensitivity or Broad Range reagents depending on estimated library concentration by Bioanalyzer analysis of libraries. Libraries were pooled and the final pool was subjected to another round of size selection with homemade Serapure beads to remove the remaining adapter dimers. Libraries were sequenced on 2 lanes of Novaseq S2 at the Chan Zuckerberg Institute Biohub.

### ChIP-seq data analysis

Reads were aligned to the Silveira genome [22] using BWA MEM 0.7.17. Peaks were called using the IP samples compared to the control input samples with macs2 version 2.2.7.1 [36], -- mfold 5-60, with option –keep-dup set to all, with the nomodel option selected, and a manually set extension size of 197 for all samples based on the estimated fragment size from Bioanalyzer traces. ChIP peaks were assigned to individual genes if any part of the peak fell in the intergenic region between the stop codon of the upstream gene and before the start codon for that gene, limiting the intergenic size to a maximum of 10kb. Subsequent gene-level analyses were performed on genes whose promoter had peaks assigned in at least 2 out of 3 replicates.

### Motif calling

Peaks that were present in all 3 replicate datasets were combined in an iterative manner into a minimal peak using the following criteria: 1) if the location of the maximum of peak 1 fell between the start and end of peak 2, and vice versa for the maximum of peak 2 falling between the start and end of peak 1. 2) A new combined peak was created using the minimal width possible by choosing 1 start and 1 end from the 2 peaks. 3) Peak 3 was then combined with this new peak using the same criteria as 1 and 2. Then, these highly reproducible peaks were compared between spherule datasets and hyphal datasets at the same timepoints (e.g. spherule day 2 and hyphal day 2) and the same overlap metric described above was used to determine if a peak was bound exclusively in spherules, exclusively in hyphae, or reproducibly in both morphologies at this timepoint. DNA sequences were extracted from these highly reproducible peak regions that fell into each of these categories and used as input for MEME [37] 5.4.1 using the flags-revcomp, -mod anr, -nmotifs 10, -w 10, -dna. Motif searches were done using MAST [38] as previously described [15].

### Generating a list of candidate transcription factors

We used the current Pfam to GO mapping from the GO Consortium (https://current.geneontology.org/ontology/external2go/pfam2go dated 2023/03/07 22:16:20) and developed a list of 246 Pfam accession numbers corresponding to GO terms containing the text “transcription factor,” “sequence-specific DNA binding,” or “regulation of DNA-templated transcription” [39, 40]. We manually added additional fungal-specific transcription factors that were not captured by GO terms (PF04082 [41], PF02292 [41], PF04769 [41], PF09729 [42], PF11754 [43, 44], PF00010 [45], PF00096 [46], PF12756 [46], PF00808 [47], PF04438 [48], PF08618 [49], PF05368 [50], PF08581 [51], and PF01722 [52]) for a total of 260 Pfam families. We then determined which genes in *Coccidioides* had these Pfam domains, based on the published genome annotation [22]. Genes with hits to 61 different transcription factor Pfam domains are present in the Silveira genome, representing 280 genes in total. This list was evaluated manually, and 53 false positives were removed for a total of 227 candidate transcription factors in *Coccidioides*.

### Hyphal radial growth assay

500 arthroconidia were resuspended in 10 µL PBS and spotted in the middle of a 2x GYE agar plate with Penicillin/Streptomycin (100 U/mL penicillin and 100 g/mL streptomycin) and incubated at 30°C. Between days 3 and 9, 3 images of the growing colonies were taken. The colony area was measured and used to calculate the radius. Hyphal growth rate is equivalent to the change in colony radius over time.

### *DitCluster*Δ mutant transmission electron microscopy

Wildtype and mutant arthroconidia were generated as described above except they were grown for 8 weeks on 2x GYE agar with Penicillin/Streptomycin (100 U/mL penicillin and 100 g/mL streptomycin) before harvest. Arthroconidia pellets were fixed in freshly prepared 2.5 % glutaraldehyde (EMS) in 0.1 M cacodylate buffer pH 7.4 (EMS) at RT for 30 minutes, then pelleted by spinning 14000 rpm for 1 minute. RT fixative was removed, and the cells were resuspended in the same fixative cooled to 4°C and stored at 4°C until ready for embedding. They were then post-fixed in 1 % OsO_4_ in 0.1 M cacodylate buffer for 1 hour on ice and then stained with 2 % uranyl acetate for 1 hour on ice. The samples were dehydrated in a graded series of ethanol washes (50 %–100 %) once, followed by a wash with 100 % ethanol and 2 washes with acetone for 15 minutes each, and then embedded with Durcupan. 70 nm sections were cut on a Leica UCT ultra-microtome and collected on 300 mesh copper grids. Sections were stained with 2 % uranyl acetate for 5 minutes, and Sato lead stain for 1 minute. Samples were viewed using a JEOL 1400-plus TEM (JEOL, Peabody, MA). Transmission electron microscopy images were taken using a Gatan OneView 4 k×4 k camera (Gatan, Pleasanton, CA).

## Results

### Transcriptomics of *Coccidioides* spherule development

Using the optimized spherulation conditions that we recently established [23], we germinated arthroconidia into spherules, observed morphology by light microscopy (Figure S1A), and isolated RNA for RNA-Seq at each timepoint from day 0 through day 6 (Figure 1A). We observed isotropic swelling in ∼25 % of arthroconidia in each replicate at 8 hours (Figure S1B). Early spherules appeared by day 1 and continued to grow until day 3 when endospore release was first observed. On days 4-6, cultures developed into a complex mixture of maturing spherules, spherules releasing endospores, free endospores, and a small proportion of hyphae (Figure 1B, example morphology Figure S1A). Each replicate at each timepoint is from an independent culture so there was some variation in the degree of endospore release in each flask (such as replicate 2, day 5). However, the overall trend is an increasing proportion of free endospores at later timepoints. As expected, spherule diameters were similar across replicates (Figure S1C). The transcriptome changed significantly over spherule development and as endospores were released, with 6355 transcripts (of 8628 total observed) undergoing at least a 2-fold change at 5 % FDR at one or more timepoints throughout this experiment (Figure 1C, Table S2). The arthroconidia stage exhibits a more divergent transcriptome than any other pairwise comparison throughout spherulation. The 8h transcriptome demonstrates moderate correlation with the arthroconidia transcriptome and a similar degree of correlation with the day 1 transcriptome (both Pearson correlation of 0.5-0.6), consistent with a transition between the arthroconidia and spherule state (Figure S1D). Day 1-6 spherules are more correlated with each other than either the arthroconidia or 8h timepoint, suggesting a consistent spherule signature. The number of transcripts that were significantly differential compared to the preceding timepoint decreased monotonically across spherulation, consistent with progression toward a more steady-state spherule transcriptome by the end of the experiment (Figure S1E). However, there are a group of transcripts that exhibit the interesting pattern of high abundance in arthroconidia and then returning to high abundance after endospores are released, suggesting these transcripts may accumulate in both spore forms (arthroconidia and endospores).

### Generating high-density transcriptomics of wildtype and mutant *Coccidioides* under spherule- and hyphal-inducing conditions

Over the course of this analysis, we used two strategies to characterize the spherule transcriptome and to identify key spherule-associated transcripts. We determined which transcripts were regulated by the critical transcriptional regulator Ryp1 (*RYP1*-dependent genes) and we compared the spherulation transcriptome to the hyphal transcriptome to identify transcripts that were associated with each morphology (morphology-dependent genes). The transcription factor Ryp1 is required for spherulation in *Coccidioides* [12]. We reasoned that understanding the portion of the spherule transcriptome that is dependent on Ryp1 would identify transcripts whose expression is associated with spherule formation rather than the conditions used to generate spherules. First, we germinated both wildtype and *ryp1*Δ arthroconidia under spherulation conditions, observed morphology by light microscopy, and performed RNA-seq at the same timepoints as the previous experiment, now sampling from the same culture over time to increase consistency between subsequent timepoints of development (Figure 2A, S2A). To compare the wildtype spherules generated in our first and second experiments, we determined the time of endospore release, the quantity of endospore release, and spherule size. The wildtype strain did release endospores starting on day 3 but there was quantitatively less endospore release in this experiment (Figure 2B). Wildtype spherules achieved a similar diameter by day 6 as seen previously (Figure S2B). Again, the arthroconidia transcriptome was the most distinct within a genotype, with sets of genes being induced/repressed as spherules formed and genes showed substantial dependence on *RYP1* (Figure 2C, Table S3). We also observed a similar pattern of decreased differential transcripts between subsequent timepoints as the experiment progressed (Figure S2C). Therefore, we conclude that these separate spherule development trajectories are comparable except for the endospore release stage.

To simultaneously query the transcriptome of the hyphal morphology, we germinated the same arthroconidia stocks of wildtype and *ryp1*Δ in hyphal-inducing conditions. It has been common to compare spherules grown in Converse medium to hyphae grown in a different rich medium (GYE) [7, 8], but, to eliminate media-specific expression effects, we generated hyphae in Converse medium (at ambient temperature without additional CO_2_). At each timepoint, we observed hyphae formation by light microscopy and the transcriptome by RNA-Seq (Figure 2D, S2D, 2E). We observed that wildtype samples formed germ tubes by day 1, with extension and early branching on day 2, followed by robust hyphal mats on day 3. We expected older hyphae to undergo arthroconidia generation and observed early evidence of arthroconidia formation on day 6 (Figure 2D open arrows). The *ryp1*Δ mutant also demonstrated rare germ tubes on day 1 but appeared to have delayed hyphal branching as we did not observe branching structures until day 3 (Figure S2D, black arrow heads). On day 6, instead of early arthroconidia development, *ryp1*Δ demonstrated aberrant morphology with chains of rounded and oblong structures (Figure 2D, black arrows), similar to the morphology at late timepoints in spherulation conditions. *ryp1*Δ arthroconidia (same biological samples as seen in Figure 2C) demonstrated significantly different expression compared to wildtype arthroconidia, but wildtype and mutant hyphal transcriptomes started to resemble each other more closely over time (Figure 2F, Table S3), suggesting that Ryp1 is largely dispensable for the hyphal transcriptome. As observed with spherulation, we observed a similar pattern of decreased differential transcripts between subsequent timepoints as the experiment progressed (Figure S2E).

### Identifying Ryp1-dependent and morphology-dependent transcripts during spherule and hyphal formation

To further refine our understanding of the *Coccidioides* transcriptome and to elucidate the molecular role of Ryp1 in *Coccidioides* development, we examined which transcripts are significantly differential in wildtype compared to the *ryp1*Δ mutant at each timepoint of spherulation or hyphal growth, termed ‘*RYP1*-dependent’ (Figure 3A). Surprisingly, the highest number of *RYP1*-dependent transcripts was in arthroconidia, where a role for Ryp1 in gene regulation has not been interrogated previously. This effect was observed regardless of arthroconidia storage conditions prior to use (Figure S3A, S3B) and indicates a previously unknown and significant role for Ryp1 in the transcriptome of arthroconidia, the infectious particle of this fungus, that bears further study.

During germination into spherules, there were increasing numbers of *RYP1*-dependent transcripts until a peak at day 4, with more transcripts induced by *RYP1* (purple) versus relatively constant numbers of transcripts repressed by *RYP1* (green) (Figure 3A). In contrast, cells in hyphal-inducing conditions trended toward fewer *RYP1*-dependent transcripts over time, indicating that the wildtype and *ryp1*Δ hyphal transcriptomes converge as both genotypes differentiated into hyphae (Figure 3A). At each timepoint sampled in both spherule and hyphal conditions, we highlighted *RYP1*-dependent transcripts that were also “morphology-dependent” (significantly differential between spherules and hyphae in wildtype culture at the same timepoint) or “morphology-independent” (Figure 3B, dark and light regions respectively). In contrast to stable numbers of *RYP1*-dependent morphology-independent genes, there was an increase in the number of *RYP1*-dependent morphology-dependent transcripts in spherules from 8 hours to day 3. This increase was largely driven by two groups of transcripts: (1) *RYP1*- activated, spherule-activated or (2) *RYP1*-repressed, hyphal-activated. This trend is more easily observed in Figure 3C, where the day 3 data are plotted, and in the global analysis in Figure S3C, S3D. In hyphal-promoting conditions, this trend was not observed, and both *RYP1*- dependent, morphology-dependent and *RYP1*-dependent morphology-independent transcripts decreased over time with no clear correlation between the hyphal-*RYP1*-dependent transcriptome and the morphology transcriptome (Figure 3B, 3C, S3C, S3D). Thus, during spherule development, *RYP1* has an impact on the morphology regulon that increases with time and peaks on day 3, as well as a morphology-independent impact with constant magnitude over time. On the other hand, in hyphal development, *RYP1* has a largely morphology-independent impact on the transcriptome that decreases over time, indicating that wildtype and *ryp1*Δ hyphae converge on similar transcriptomes.

**Figure 3:**
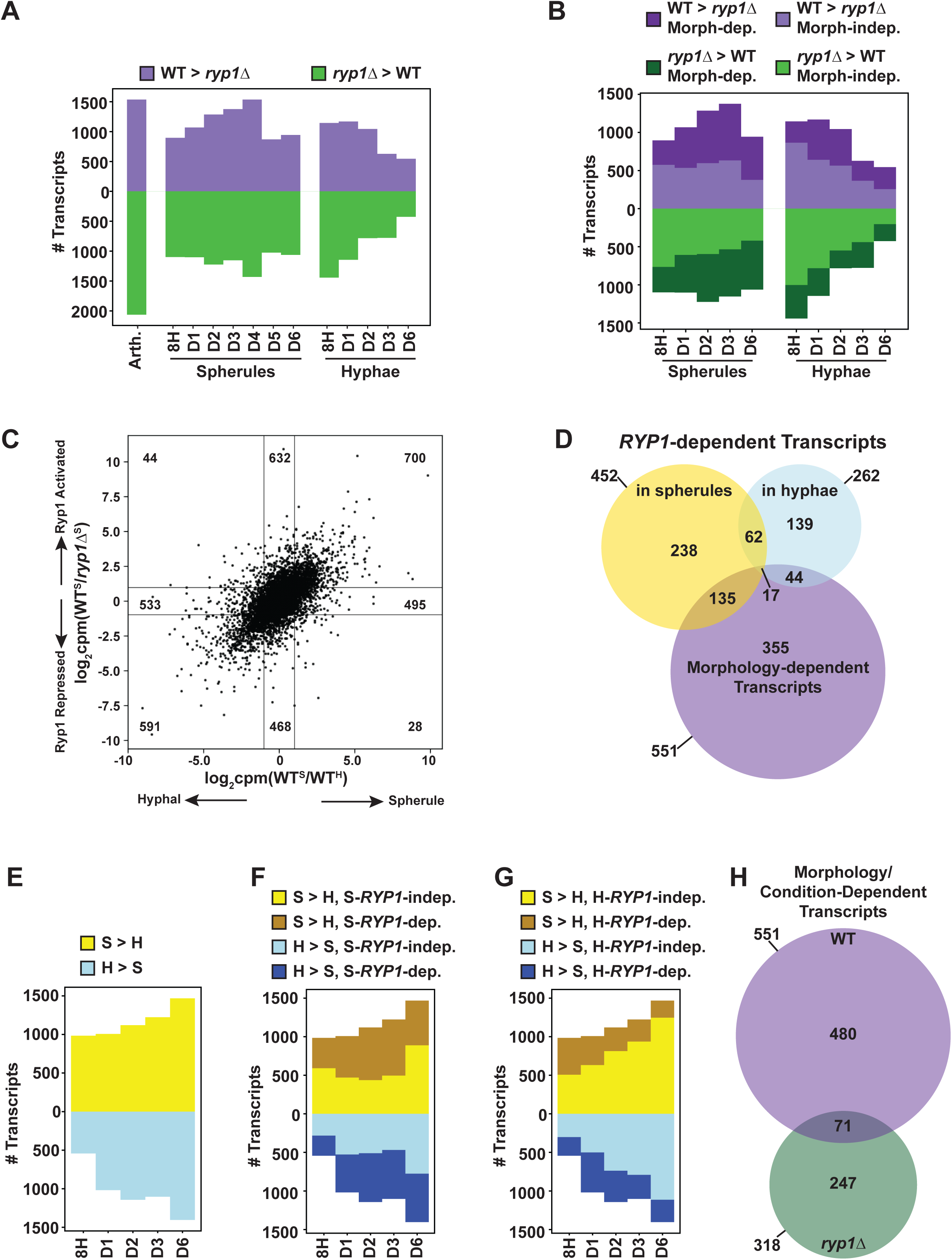
Defining *RYP1*-dependent and morphology-dependent transcripts. A. Number of significantly-differential transcripts between wildtype and the *ryp1*Δ mutant at each timepoint specified. Transcripts that are induced by *RYP1* (higher in wildtype than *ryp1*Δ) are in purple and transcripts that are repressed by *RYP1* (higher in *ryp1*Δ than wildtype) are in green. B. As in A but only with timepoints where paired spherule and hyphal wildtype datasets are available to highlight morphology-dependent genes. Dark purple and dark green correspond to the number of *RYP1*-dependent transcripts that are also morphology-dependent (significantly differential between wildtype spherules and wildtype hyphae) at that timepoint. C. Scatterplot demonstrating expression of all detected transcripts at day 3. On the x-axis, values are the ratio of wildtype spherule over wildtype hyphal transcript abundance transformed to log_2_(counts per million). On the y-axis, values are the ratio of transcript abundance of wildtype spherule over *ryp1*Δ in spherulation conditions, transformed to log_2_(counts per million). D. Overlap between transcripts that are significantly differential between wildtype and the *ryp1*Δ mutant at all timepoints of spherulation (day 1-6, excluding the 8h timepoints since spherules and hyphae had not appeared by then), transcripts that are differentially expressed between wildtype and the *ryp1*Δ mutant over all timepoints of hyphal formation (day 1, 2, 3, 6), and transcripts that are morphology-dependent in wildtype at all comparable timepoints (day 1, 2, 3, 6). E. Number of significantly differential transcripts between wildtype spherules and hyphae at each timepoint specified. Transcripts with higher abundance in spherules than hyphae are yellow and transcripts with higher abundance in hyphae than spherules are blue. F. As in E, now highlighting dark yellow and dark blue transcripts corresponding to the number of morphology-dependent transcripts that are also regulated by *RYP1* at the corresponding spherule timepoint. G. Same graph as F except the dark yellow and dark blue transcripts now correspond to the number of morphology-dependent transcripts that are regulated by *RYP1* at the corresponding hyphal timepoint. H. Overlap between transcripts that are significantly differential between wildtype spherules and hyphae at all comparable timepoints (day 1, 2, 3, 6) and transcripts that are differentially expressed between the *ryp1*Δ mutant in spherulation and hyphal-inducing conditions at the same timepoints.

Next, to further understand the role of Ryp1 in *Coccidioides* biology, we examined the stringent set of transcripts that were *RYP1*-dependent across all spherule or hyphal timepoints (Figure 3D). There were 452 transcripts consistently *RYP1*-dependent across all 6 timepoints in spherulation conditions (termed “S-*RYP1*-dependent”) and 262 transcripts across all 4 timepoints in hyphal conditions (termed “H-*RYP1*-dependent”). Of these 452 S-*RYP1*-dependent and 262 H-*RYP1*-dependent genes, 79 were common to both sets (p = 1.94e-38 by Fisher Exact test). While significant, this relatively low magnitude of overlap adds additional evidence for *RYP1*’s distinct regulatory roles in spherule and hyphae. We also found 551 consistently morphology-dependent transcripts by comparing wildtype spherules and hyphae. 152 (135 + 17) of these strictly morphology-dependent transcripts were also consistently S-*RYP1*- dependent (p = 4.97e-71 by Fisher Exact test) and 61 (44 + 17) strictly morphology-dependent genes were consistently H-*RYP1*-dependent (p = 1.69e-18 by Fisher Exact test). The 17 transcripts that are S-*RYP1*-dependent, H-*RYP1*-dependent, and morphology-dependent include the gene D8B26_005342/CIMG_00509, which is already known to be spherule-induced, *RYP1*-dependent [12], and interestingly in an area of genomic introgression between the two known *Coccidioides* species, *C. posadasii* and *C. immitis* [53]. This central overlap of 17 is surprisingly low although still significantly higher than expected by chance (χ^2^ = 1019.27, p < 0.0001) and again implies that *RYP1* has 2 distinct regulatory roles in these two morphologies, more significant in spherules compared to hyphae. The high number of morphology-dependent genes that are *RYP1*-independent suggests roles for additional regulators of spherulation.

### Morphology change triggers differential expression of a core set of transcripts across all developmental timepoints

Next, we examined morphology-dependent transcripts at each shared timepoint of spherule and hyphal development in wildtype (Figure 3E). As expected, the number of morphology-dependent transcripts increased over time as spherules and hyphae emerged. We highlighted the morphology-dependent transcripts that were also S-*RYP1*-dependent (Figure 3F) at the same timepoints and observed an increase in the magnitude of this subset of transcripts with the exception of day 6, while morphology-dependent S-*RYP1*-independent genes remain relatively constant (with the same exception of day 6). On the other hand, when we highlighted the number of morphology-dependent-genes that are H-*RYP1*-dependent at the same timepoints, that number is relatively small (Figure 3G) and does not have a clear trend. Thus, the role of *RYP1* in regulating morphology is related to its regulon in spherulation conditions, where it induces spherule-associated transcripts and suppresses hyphal-associated transcripts. In hyphae, *RYP1* regulates a small subset of transcripts but most of these seem to be morphology-independent.

Finally, we defined a stringent set of transcripts that were consistently morphology-dependent across all shared spherule and hyphal timepoints (Figure 3H). As discussed above, 551 transcripts were consistently morphology-dependent in wildtype. 318 transcripts were consistently differential in the *ryp1*Δ mutant growing in spherulation conditions compared to hyphal conditions, even though the mutant forms hyphae under both these conditions. Given the uniform morphology, these 318 transcripts are likely responding to the difference in spherulation- and hyphal-inducing conditions (namely, temperature and CO_2_). Surprisingly, the overlap between the 551 morphology-dependent genes in wildtype and the 318 condition-dependent transcripts in *ryp1*Δ is low in magnitude (71 transcripts total, p = 2.56e-20 by Fisher Exact test), meaning that the majority of the 551 morphology-dependent transcripts are linked to the morphology itself.

Focusing on the 551 transcripts with morphology-dependent expression in wildtype, 273 are consistently spherule enriched (of those, 82 are also consistently S-*RYP1*-dependent) and 239 are hyphal enriched (of those, 32 are also consistently H-*RYP1*-dependent). We examined these subsets further at the gene level to better understand the molecules involved in the *Coccidioides* morphologic transition. Within the spherule-enriched set, as expected, we found the transcript encoding the best characterized virulence factor in *Coccidioides*, SOWgp [54] (D8B26_003939) and the previously-reported spherule-associated gene *PSP1* [7, 8, 12, 55] (D8B26_002733). We also found D8B26_003869, the ortholog of *BOI2* in *Saccharomyces cerevisiae*, a gene involved in polar growth and inhibition of cytokinesis during budding [56], which may imply a role for directed vesicle fusion with the plasma membrane or a delay in cytokinesis during spherule development. Additionally, there are two transcription factors (D8B26_005038 and D8B26_006698) in this group that are good candidates for regulators of spherulation in addition to Ryp1. Of note, *OPS1* [12, 55] (D8B26_004398) and *ALD1* [55, 57] (D8B26_007314), genes that were previously published to be spherule-biased, were found to be spherule enriched in some early timepoints but not consistently at later timepoints of morphological development, demonstrating the power of this high-density developmental time course. Finally, despite the critical role Ryp1 plays in inducing spherulation in *Coccidioides*, the *RYP1* transcript itself does not demonstrate morphology-specific expression (Figure S3E). In the consistently hyphal-enriched transcripts, we found *STU1* (D8B26_002234), the *Coccidioides* ortholog of *Aspergillus* APSES family transcription factor *STUA* which regulates conidiation [58]. This finding matches the ortholog of *STUA* in *Histoplasma*, *STU1/EFG1*, which is extremely hyphal-biased in its expression [59]. Consistent with previous findings, the major component of the woronin body structure that plugs damaged areas of hyphal walls, *HEX1* (D8B26_006047), was upregulated in hyphal conditions compared to spherules [7]. As expected, these hyphal-associated genes were also upregulated in *ryp1*Δ cells in both spherule- and hyphal-inducing conditions. Somewhat unexpectedly, the cytosolic catalase (D8B26_007217) was found to be consistently higher in hyphal conditions and the *ryp1*Δ mutant. This gene has been previously found to have higher expression in spherules than hyphae [8, 12] in studies in which the spherules and hyphae were grown in different media. Given the discordant findings between our data and previous publications, we believe nutritional cues play a key role in regulating this particular transcript. Thus, our rich dataset identifies 551 consistently morphology-dependent transcripts that are prime effector and regulatory candidates for control of the *Coccidioides* developmental program, deconvolutes the effects of change in growth conditions from change in morphology, and identifies 273 spherule-enriched genes that are likely to be involved in virulence.

### Ryp1 binds to two distinct subsets of promoters

We next sought to determine which *RYP1*-dependent genes displayed association with Ryp1 using ChIP-Seq with an antibody generated against a peptide epitope of Ryp1. While we attempted to perform ChIP on arthroconidia and multiple timepoints of spherule or hyphal growth (8h, D1, D2, D4, micrographs in Figure S4A) and one timepoint for each morphology of the *ryp1*Δ mutant (micrographs in Figure S4B), consistent Ryp1 binding was only detectable for spherules on days 1, 2, and 4 and hyphae on days 2 and 4 (Table S4). This lack of binding in wildtype may be due to less initial biomass and does not necessarily reflect a lack of Ryp1 binding at those early timepoints. In the *ryp1*Δ mutant, we expected very little binding of the Ryp1 antibody and, while we did identify sporadic peaks in individual replicates, they were not reproducible and likely represented low-level off-target binding of the antibody. We focused our subsequent analyses on those later timepoints with >400 detected peaks in at least 2 of 3 replicates. As expected, we observed Ryp1 binding in spherules at the *SOWgp* promoter (Figure 4A), consistent with observations in this and prior studies [11, 12, 60] that have found *SOWgp* expression to be *RYP1*-dependent. Additionally, we examined the *RYP1* locus itself and found that Ryp1 bound both upstream and downstream of the gene, suggesting a possible autoregulatory mechanism for *RYP1*, a known characteristic for Ryp1 orthologs in other fungi [15, 16, 61, 62] (Figure 4B). Finally, we found examples of Ryp1 binding in hyphae and spherules (Figure 4C), including D8B26_005360 and D8B26_005361, both *RYP1*-repressed transcripts that encode hypothetical proteins. Upon manually reviewing the 32 genes designated as bound in hyphal samples only, we found evidence of binding in spherule samples as well and believe these are instances in which MACS did not correctly identify peaks in the paired spherule timepoint. Therefore, we do not think there are any examples of Ryp1 binding promoters in hyphal samples alone.

**Figure 4:**
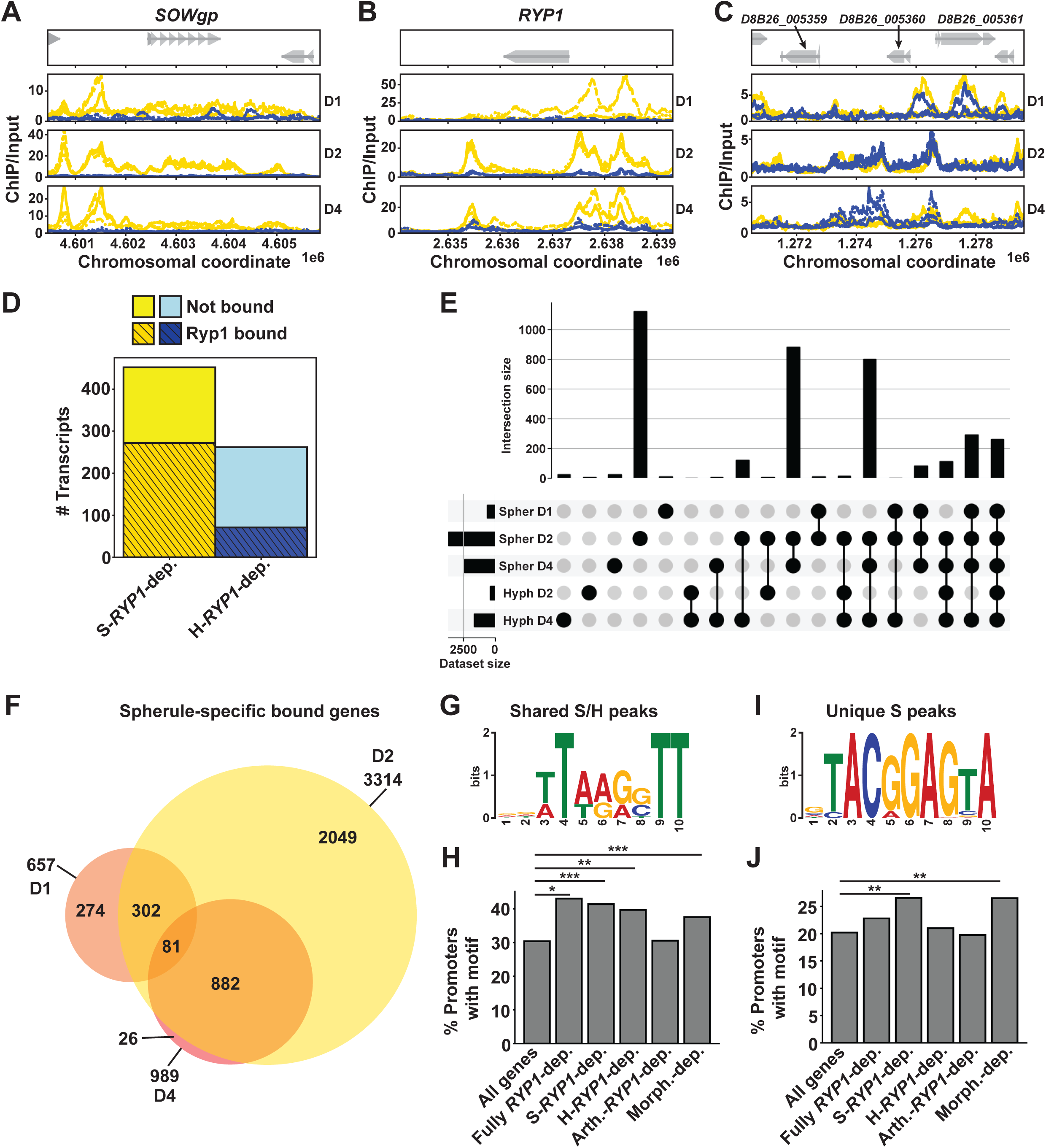
Ryp1 regulates two distinct subsets of targets. A. Traces demonstrating chromosomal location of fold enrichment of ChIP signal/input in spherules (yellow) and hyphae (blue) at the designated timepoints relative to the annotated *SOWgp* gene. B. As in A but demonstrating ChIP signal/input relative to the annotated *RYP1* gene. C. As in A but demonstrating ChIP signal/input relative to *D8B26_005359* and *D8B26_005360* genes (genes indicated by grey arrows). D. Barplot demonstrating the proportion of S-*RYP1*-dependent genes or H-*RYP1*-dependent genes (as defined in 3D) whose promoters have Ryp1 binding by ChIP- seq. E. UpSet plot demonstrating the size of each individual set of genes whose promoters are bound at each designated spherule/hyphal timepoint (bottom left) and size of overlap between each of these sets (magnitude on top, overlapping sets demonstrated by connected black circles on bottom). Each gene can only be assigned to one unique category. F. Overlap of spherule-specific peaks (genes whose promoter is bound in spherule timepoints only, without Ryp1 binding in the corresponding hyphal timepoint or *ryp1*Δ mutant subjected to the same conditions as wildtype) on days 1, 2, and 4. G. Motif enriched in DNA sequences of Ryp1 ChIP- seq peaks found consistently in both day 2 and 4 spherule and hyphal datasets (498 sites p = 4.7e-064 on day 4 and 193 sites p = 3.7e-046 on day 2). H. Percent of genes in each subset (from 3A, 3D) whose promoter regions have at least one hit for the Ryp1 binding motif in 4G. Promoters are defined as the sequence upstream of the CDS start until the next upstream CDS is encountered, or 10kb maximum. Fully *RYP1*-dependent genes are those that are significantly differential between wildtype and *ryp1*Δ in all spherule timepoints (day 1-6) and all hyphal timepoints (day 1, 2, 3, 6). *: p <0.05, **: p <0.005, by Fisher exact test. I. Motif enriched in DNA sequences of Ryp1 ChIP-seq peaks found uniquely in day 2 and 4 spherule datasets (>1000 sites p = 1.3e-044 on day 2, 215 sites p = 4.5e-046 on day 4). J. As in H but now for Ryp1 binding motif in I.

We determined which *RYP1*-dependent genes observed by RNA-seq were also direct targets of Ryp1 by ChIP-Seq. We found that 60 % of the 452 S-*RYP1*-dependent transcripts also demonstrated Ryp1 promoter binding in at least one spherule timepoint of our ChIP-seq experiment (Figure 4D). In contrast, only 27 % of the 262 H-*RYP1*-dependent genes had Ryp1 promoter binding in hyphae. We additionally examined the percentage of the 551 morphology-dependent genes we had previously defined (and the 786 morphology-dependent genes we found from the paired RNA-Seq from this ChIP-Seq experiment, Table S5) that had Ryp1 binding in their promoters. We found ∼55 % of both subsets of morphology-dependent genes were Ryp1 targets (Figure S4C). Thus, Ryp1 plays an important role as a direct regulator of morphology in *Coccidioides*, where it seems to act specifically by binding promoters in spherule development. Although the loss of *RYP1* influences the hyphal transcriptome, the ChIP-Seq data suggest that effect is more indirect.

Next, we created an UpSet plot (Figure 4E) to group genes whose promoters had Ryp1 binding detected. This analysis revealed that most genes fell into 3 categories: 1) genes whose promoters are bound at the spherule day 2 timepoint only, 2) genes whose promoters are bound in both later spherule timepoints, and 3) genes whose promoters are bound in both later spherule timepoints in addition to the latest hyphal timepoint. Taken together, this likely indicates 2 distinct regulons for Ryp1: an exclusive spherule regulon and a shared spherule/hyphal regulon. Of note, there were minimal numbers of genes whose promoters demonstrate Ryp1 binding in hyphae only, indicating that there does not seem to be a unique Ryp1 regulon in hyphae. Given the large number of spherule Ryp1 peaks detected, we further delved into those genes that had spherule-specific promoter binding (without any observed Ryp1 peaks in hyphal samples or *ryp1*Δ samples). In examining the genes containing spherule-specific peaks across the spherule timepoints (Figure 4F), we found that the Ryp1 spherule regulon appears to be dynamic, with 274 genes bound by Ryp1 only in day 1 spherules, 2049 unique genes bound by Ryp1 in day 2 spherules, and 882 genes bound by Ryp1 in both of the later spherule timepoints. Thus, Ryp1 in *Coccidioides* appears to have a complex and dynamic role in regulating multiple stages of the spherule morphology, suggesting it plays a role in a complex regulatory network like some of its orthologs in other fungi [15, 63, 64].

### Two distinct *RYP1* motifs in *Coccidioides* are enriched in promoters of spherule *RYP1*- dependent genes

To better understand how Ryp1 can regulate two distinct subsets of genes, we performed motif searches on multiple subsets of Ryp1 peaks combined as illustrated in Figure S4D: (1) peaks found in promoters of genes in both spherules and hyphae and (2) peaks found in promoters of genes only in spherules. In the first group, we discovered a significantly-enriched motif (Figure 4G) that is extremely similar to the previously published Ryp1 motif in *Histoplasma* (Figure S4E). We used MAST to search for this Ryp1 spherule/hyphal motif in all *Coccidioides* promoters in the genome with the threshold E-value of 2.08e-04, a cutoff which proved useful for this analysis in *Histoplasma* [15]. Since the motif has low information content, we found 30 % of all promoters had a hit to the Ryp1 motif (Figure 4H). The percent of promoters containing Ryp1 motif hits was highest for genes whose transcripts are *RYP1*-dependent across all timepoints studied and genes whose transcripts are *RYP1*-dependent across all spherule timepoints (S-*RYP1*-dependent). Since this motif was derived from peaks found in both spherule and hyphal morphologies, unsurprisingly, the enrichment of the Ryp1 binding motif in the promoters of H-*RYP1*-dependent genes and morphology-dependent genes was also quite high (∼40 %). Finally, we found no enrichment of the motif in the 3599 *RYP1*-dependent transcripts in arthroconidia, suggesting that *RYP1* may control this regulon indirectly through a second major regulator. Together, this motif analysis indicates that the presence of the Ryp1 motif in promoters alone does not explain the varying impact of *RYP1* on spherules, hyphae, and arthroconidia that we observed by RNA-seq. Next, we examined the number of Ryp1 motif hits per promoter for each of these subsets of Ryp1-motif-containing promoters (Figure S4F). Interestingly, about 40 % of S-*RYP1*-dependent genes with motif hits had more than 1 motif hit, including some promoters with up to 16 total Ryp1 motif hits. This trend toward more motif hits was unique to S-*RYP1*-depenent genes and may provide a clue toward the mechanism by which Ryp1 has more impact on the transcriptome in spherules despite a shared DNA binding sequence in both spherules and hyphae.

Second, we performed motif searches on peaks that were found only in spherule conditions and discovered a novel motif that has not been reported before for Ryp1 association in any organism (Figure 4I). Given the extremely different sequence from the canonical Ryp1 motif described above, we hypothesize that this motif reflects recruitment of Ryp1 to these promoters via interaction with a second (unknown) regulator that binds this motif directly. Using a more stringent E-value of 1e-06 given the higher information content in this motif, we found that it was present in 20% of promoters across the genome. This motif was significantly enriched in S-*RYP1*-dependent and morphology-dependent gene promoters (Figure 4J). Unlike the canonical Ryp1 motif discussed above, there was not a similar trend toward increased numbers of Ryp1 motif hits per promoter in any gene subsets (Figure S4G). Interestingly, S- *RYP1*-dependent and morphology-dependent genes have significantly longer promoters than other gene subsets, which may accommodate both motifs we identified and potentially more numbers of the canonical Ryp1 motif in S-*RYP1*-dependent genes (Figure S4H). Thus, we find that distinct Ryp1-associated motifs, number of motifs per promoter, and potentially combinatorial motifs could contribute to the ability of Ryp1 to possess 2 distinct regulons.

### Candidate transcription factors for regulation of spherulation

As the effect of Ryp1 on the transcriptome does not fully explain the morphology transition of *Coccidioides*, there are likely additional regulators involved in switching morphology and the maintenance of spherulation. To generate additional candidates, we created a list of 227 possible transcription factors in *Coccidioides* and examined their expression in the RNA-Seq data from the experiments in Figure 2 (Figure 5A, Table S6). Most of the transcription factor candidates are expressed more highly in spherules compared to hyphae, including *RYP2* and *RYP4* which are part of a regulatory network that acts with *RYP1* to control morphology of the related fungus *Histoplasma* [15]. Interestingly, a cluster of 46 transcription factors that are highly expressed in late spherule timepoints are also highly expressed in arthroconidia, including *RYP4, PAC2* [65] (the paralog of *RYP1*), and *VEA1* (a velvet protein like *RYP2* and *RYP3).* Velvet proteins are unique to fungi and function in regulating developmental processes and secondary metabolism [66]. Transcription factors that are expressed in hyphae more than spherules include *STU1* and *FBC1,* both known to be expressed in hyphae in *Histoplasma* [67, 68] and involved in hyphal development and conidiogenesis in *Aspergillus nidulans* [69, 70]. Excluding *RYP1* itself, 16 transcription factor candidates were found in the 452 S-*RYP1*- dependent transcripts defined above, including 3 candidates that were also found in the 262 H- *RYP1*-dependent transcripts. 14 of the 16 S-*RYP1*-dependent transcription factor candidates also exhibited Ryp1 binding in at least 1 spherule timepoint. Given the central role of *RYP1* in regulating the morphology transition, these 14 transcription factors that are direct regulatory targets of *RYP1* are good candidates to be additional members of the regulatory network that controls morphology in *Coccidioides*. 10 of these candidates are repressed by *RYP1* in spherules and include *FBC1*. The remainder of these direct *RYP1* targets, including the 4 that are induced by *RYP1* in spherules, are currently unannotated.

**Figure 5:**
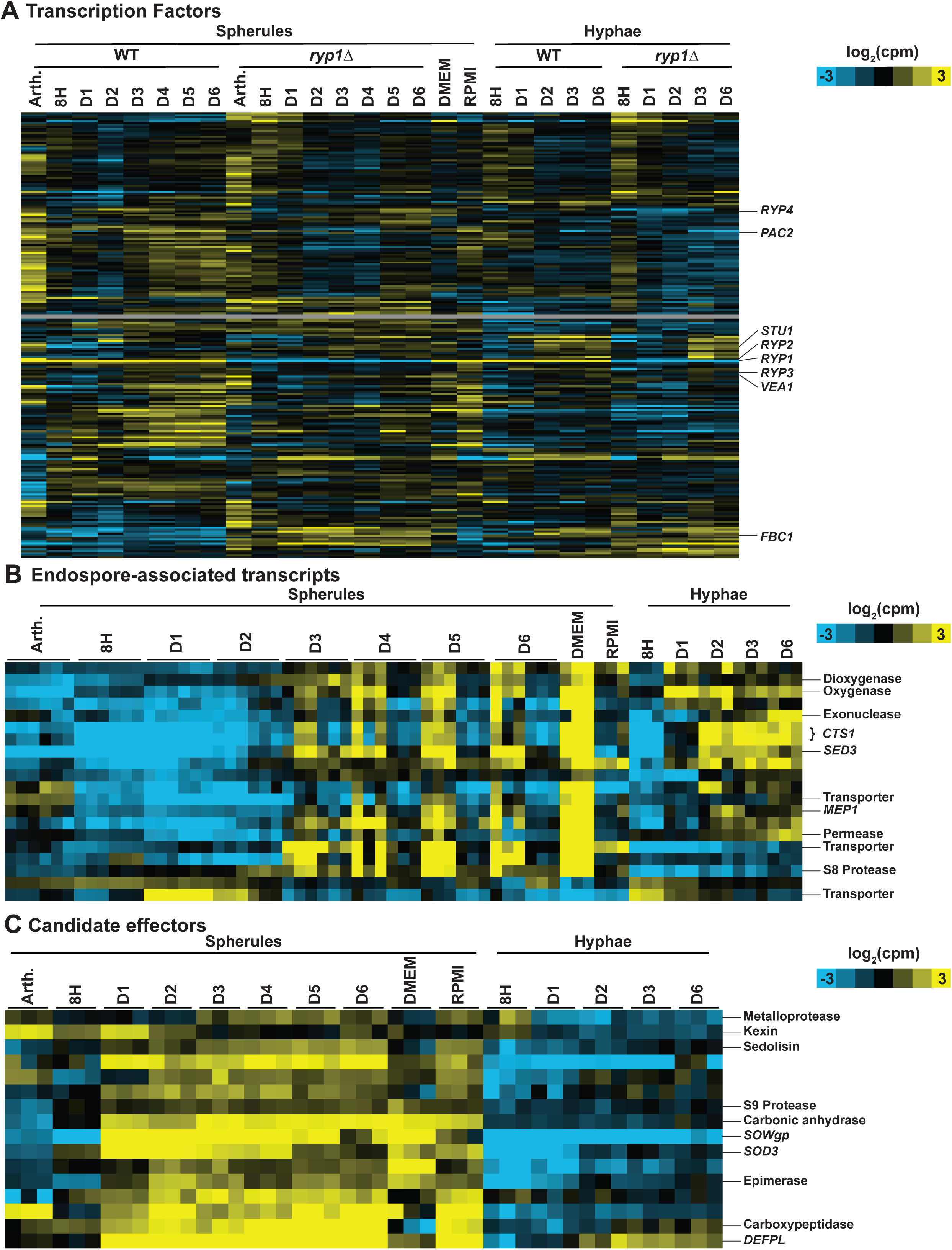
Defining *Coccidioides* transcription factors, endospore-associated genes, and candidate effectors over spherulation and hyphal development. A. Heatmap of transcript abundance for all predicted transcription factors in *Coccidioides*. Expression data all from experiments described in Figure 2, with additional DMEM and RPMI conditions as described in S5A, S5B. Rows are clustered based on correlation across all columns. Log_2_(counts per million) indicated by yellow and blue shading. Grey rows indicate transcription factors that did not have sufficient counts to pass the filter threshold for RNA-Seq analysis. B. Heatmap of transcript abundance for endospore-associated genes. Each spherule timepoint shows 6 replicates: the first 3 are from the experiment in Figure 1 and the last 3 are from the experiment in Figure 2. DMEM and RPMI timepoints are as described in Figure S5A, S5B. Rows are clustered based on correlation across all columns. Log_2_(counts per million) indicated by yellow and blue shading. C. Heatmap of transcript abundance for putative spherule-expressed secreted effectors. Expression data all from experiment described in Figure 2, with additional DMEM and RPMI conditions as described in S5A, S5B. Rows are clustered based on correlation across all columns. Log_2_(counts per million) indicated by yellow and blue shading.

### Defining endospore-associated transcripts

Since the endospore form is even less characterized than spherules, we used the RNA-seq data corresponding to cultures for which we observed the most endospore release to identify potential endospore-enriched transcripts. Specifically, we interrogated day 3 and later timepoints from Figure 1, and also performed additional RNA-seq from spherules formed in DMEM + 20 % FBS and harvested at day 3, when released endospores were abundant (Figure S5A). We defined endospore-enriched transcripts as those that were all consistently differential on days 3- 6 compared to days 1 and 2 in the experiment from Figure 1, all significantly differential in DMEM conditions compared to RPMI + 10 % FBS conditions (Figure S5B) which did not exhibit endospore release, and all significantly differential in DMEM conditions compared to day 1 and 2 spherules, day 1 and 2 hyphae, and arthroconidia from the same experiment. For all these differential comparisons, we enforced criteria that the direction of differential expression had to be consistent for the transcript across the comparisons made. Of the transcripts meeting the above criteria, there were 18 transcripts with an increase in expression in samples containing endospores and 2 transcripts that demonstrated consistent decrease in expression in samples containing endospores (Figure 5B, Table S7). The transcripts with increased abundance included *MEP1*, a metalloprotease which is known to play a role in masking endospore recognition by the immune system [71], and *CTS1* (misannotated as 2 separate transcripts D8B26_000666/7 in the current genome), an endochitinase that has been previously characterized to have maximal expression when endospores are present in culture [72]. Interestingly, the endospore-enriched transcripts include 2 other secreted serine proteases, D8B26_003356 and D8B26_007338. Both transcripts that are consistently downregulated in endospore-containing cultures have no available annotation data. This list of genes represents the first and strongest candidates for factors intimately involved in endospore biology.

### *Coccidioides* spherule secreted effectors are enriched for proteases

Next, we set out to define a set of putative secreted effectors in *Coccidioides*, since these factors are top candidates for interaction with the host immune system. We extrapolated likely characteristics from plant fungal pathogens, where secreted effectors have been extensively studied [73, 74]. We filtered for transcripts containing a predicted signal sequence (SignalP 6.0 [75]), that are cysteine-rich (predicted protein product contains ≥ 4 cysteines), and whose expression is consistently higher in spherules than hyphae at all timepoints. This yielded a list of 16 genes, 4 of which were also *RYP1*-dependent at all spherule timepoints (Figure 5C, Table S8): D8B26_003939 which encodes SOWgp, the major component of the spherule outer wall that is exclusively expressed in the spherule form; *DEFPL* (D8B26_005342) which encodes a cysteine-rich protein of unknown function previously published to be highly upregulated in the spherule morphology [24]; D8B26_005613 which encodes a 245-aa protein of unknown function; and D8B26_005065, a serine carboxypeptidase. Surprisingly, the remaining 12 putative secreted effectors also include 5 additional proteases (a kexin, an M35 metalloprotease, another serine carboxypeptidase, an S8-like protease, and an S9-prolyl-peptidase). Therefore, more than 40 % of the putative secreted effectors we define are secreted proteases and suggest a possible protease-based virulence strategy for *Coccidioides*.

### A cluster of 6 genes that demonstrate spore-associated *RYP1*-dependent expression affects arthroconidia cell wall development

Finally, to demonstrate the ability of this extensive resource to uncover new biology, we synthesized the above data and focused on a cluster of 6 adjacent genes (D8B26_005432 to D8B26_005438, Figure 6A) which demonstrate high transcript accumulation in both arthroconidia and endospores, the 2 spore forms characterized in this study (Figure 6B). This 6- gene cluster includes *DIT1* (D8B26_005435) and *DIT2* (D8B26_005434) [76], genes encoding enzymes predicted to synthesize dityrosine, and *DTR1* (misannotated as D8B26_005432/3 in the current genome) [77], which encodes a bisformyl dityrosine transporter. In *Saccharomyces cerevisiae*, the orthologs of these genes are involved in the synthesis and assembly of the protective dityrosine layer of *S. cerevisiae* ascospores [78], suggesting the hypothesis that dityrosine or a derivative thereof may play an important role in *Coccidioides* arthroconidia and endospores as well. Given the major role for *RYP1* in arthroconidia biology that we determined from our transcriptomics, we examined whether this cluster of spore-associated genes was regulated by *RYP1*. In *Coccidioides*, the transcripts from this cluster demonstrate a complex dependence on *RYP1* (Table S9). All members of the cluster are significantly *RYP1*-dependent in at least two timepoints of spherulation and there is a trend for members of the cluster to require *RYP1* for increased transcript abundance later in spherulation, consistent with three members of the cluster exhibiting Ryp1 binding in later spherule timepoints by ChIP-Seq (Figure 6C). We interpreted these *RYP1*-dependent results as additional support for this cluster playing a role in *Coccidioides* spore biology. Therefore, we interrogated the biological role of this cluster by creating two independent deletion mutants lacking all 6 genes in the cluster (*DitClusterΔ-* 1&2) (Figure 6A). We generated arthroconidia from these mutants and wildtype and examined them by TEM. We found that mutant arthroconidia exhibited thinner cell walls than wildtype (Figure 6D, 6E). Correlated with this, we also found a modest decrease in the number of visible cell wall layers in the mutant arthroconidia as well (Figure S6A). Wildtype arthroconidia in our strain background also produce a yellow pigment that is spore-associated. In the mutant lacking the full cluster of 6 genes, the yellow color is decreased. To further interrogate this, we also created two independent deletion mutants lacking only *DIT1*, *DIT2*, and *DTR1* (*DitClusterSmall*Δ-1&2) and found that these mutant arthroconidia demonstrate a similar yellow color as wildtype (Figure S6B, S6C). This suggests that the three unnamed genes in this cluster play a role in producing the yellow pigment. As expected, none of the mutants had defects in hyphal growth or spherule formation (Figure S6D, S6E, S6F, S6G). Thus, the criteria we applied to our rich transcriptomic and ChIP-Seq atlas of *Coccidioides* development correctly predicted a role for these genes in a specific developmental morphology of *Coccidioides*. We anticipate this resource can be used in a similar manner for many additional targets, enabling a much deeper understanding of the biology of this important fungal pathogen.

**Figure 6:**
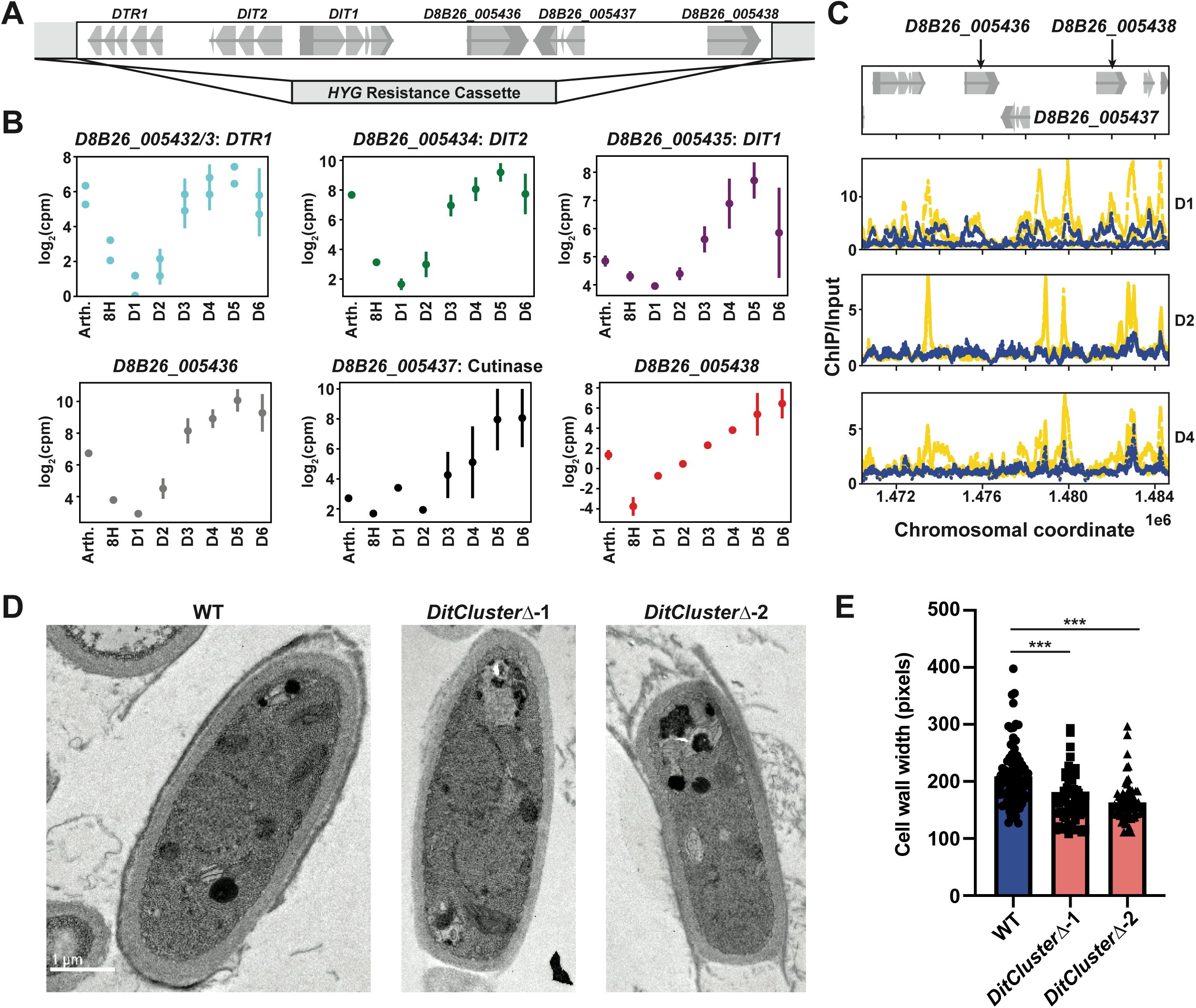
A six-gene cluster plays a role in spore development. A. Schematic of the location of the 6 genes in the spore-related cluster and position of hygromycin cassette integration. B. Plots of log_2_(counts per million) over spherulation from data in Figure 1 for each gene in the cluster. *DIT1*/*DIT2*/*DTR1* names from *S. cerevisiae* orthologs and ‘Cutinase’ name based on Pfam hit to that domain. C. Traces demonstrating chromosomal location of fold enrichment of ChIP signal/input in spherules (yellow) and hyphae (blue) at the designated timepoints relative to *D8B26_005436* - *D8B26_005438* genes. D. Representative TEM images of wildtype and each *DitCluster*Δ mutant. E. Quantification of arthroconidia cell wall width measurements of TEM images for wildtype and each *DitCluster*Δ mutant. ***: p <0.0001, by unpaired t-test.

## Discussion

We report the first transcriptomic atlas of *Coccidioides* developmental programs from vegetative arthroconidia into spherules releasing endospores, and from arthroconidia into mature hyphal mats, with almost every transcript in the *Coccidioides* genome demonstrating significant differential expression over the conditions we interrogated. These developmental programs triggered a near-full remodeling of the transcriptome. By characterizing the regulatory targets of the major morphologic regulator Ryp1 by ChIP-Seq in *Coccidioides* for the first time, our work demonstrates a clear and specific role for Ryp1 in spherules, a significant but indirect role regulating the transcriptome in arthroconidia, and a shared morphology-independent regulatory role for some spherule and hyphal genes. Using this transcriptomic atlas, we define 20 endospore-associated genes and 16 putative secreted effectors, 6 of which are, remarkably, all secreted proteases.

### Spherulation is a developmental program

Our data show that spherule development requires near-complete remodeling of the transcriptome, a level of complexity on par or surpassing developmental trajectories of multicellular organisms [79–81]. In fact, the spherule form should be considered a multicellular morphology since it is filled with hundreds of endospores which each contain one or more of their own nuclei. While elegant observational work has established the spherulation cycle and associated morphologies, we have much more to learn about this cell type. We present bulk RNA-seq here as the first step toward understanding the spherule. However, we acknowledge that there could be (and likely is based on precedent from studies of spores in other organisms) significant heterogeneity between the transcriptome of individual endospores that comprise mature spherules. Additionally, it is unknown whether the spherule retains one or more nuclei that do not develop into endospores and whether there is still a spherule-specific cytoplasm surrounding endospores that may carry out different biologic functions from the endospores themselves. The application of single cell RNA-seq and spatial transcriptomics may allow further insights into this aspect of spherulation.

### Ryp1 plays a major regulatory role in arthroconidia

We report the first transcriptomes of arthroconidia in *Coccidioides*, which are completely distinct from the spherule and hyphal transcriptomes. Surprisingly, using the metric of number of transcripts that are significantly differential in abundance between wildtype and *ryp1Δ, RYP1* plays a much larger role in regulation of the arthroconidia transcriptome than in spherules or hyphae. We were not able to detect Ryp1 binding by ChIP-Seq in arthroconidia, likely due to too little starting material. However, we used the Ryp1 binding motifs defined in this study, searched for them in promoters of *RYP1*-regulated genes in arthroconidia as defined by RNA-Seq, and found no significant enrichment. These data strongly suggest that Ryp1 impacts a second regulator that directly binds the DNA via a distinct motif, or that a co-regulator modulates Ryp1 binding for this subset of genes. Since the *ryp1*Δ mutant has decreased arthroconidia viability [12] and there is precedent that *RYP1/WOR1* is required for proper regulation of conidial development in other fungi [20, 63, 82], we interpret our findings as indicating that Ryp1 likely is a major but indirect regulator in *Coccidioides* arthroconidia as well.

### Ryp1 plays a more specific role regulating expression in spherules than in hyphae

While the *ryp1*Δ transcriptome significantly differs from wildtype in arthroconidia and hyphae as well as spherules, our analyses of Ryp1 binding by ChIP-Seq indicates that S-*RYP1*-dependent genes defined by RNA-Seq are much more likely to be direct regulatory targets of Ryp1 than H- *RYP1*-dependent genes. Additionally, there is a large set of genes that only exhibit Ryp1 binding in the spherule morphology and essentially no genes that exhibit Ryp1 binding exclusively in the hyphal morphology. Therefore, our data support a model in which Ryp1 regulates 4 distinct gene subsets: (1) a core set of morphology-independent genes in both spherules and hyphae, (2) a morphology-specific set of genes in spherules that it modulates through direct association with their promoters, (3) a set of hyphal genes whose RNA level is modulated by *RYP1* but likely through another regulator as Ryp1 does not directly associate with their promoters, and (4) a large set of genes in arthroconidia, again likely through indirect effects through another regulator (Figure 7).

**Figure 7:**
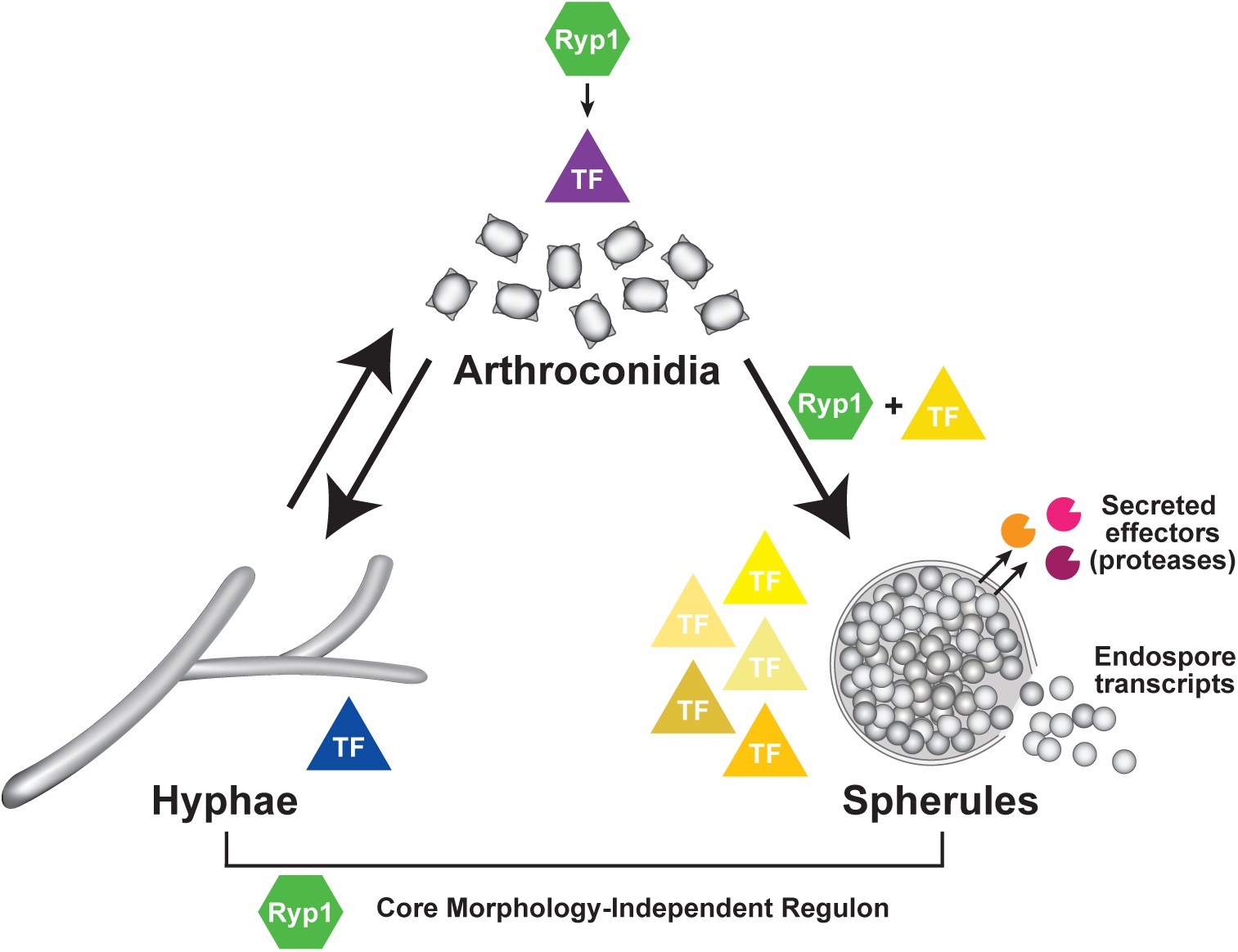
Regulation of gene expression in *Coccidioides* development. Our data uncover regulatory modules in *Coccidioides* development. The majority of *Coccidioides* transcription factors (TFs), depicted as triangles in the figure, show enhanced expression in spherules compared to hyphae. These TFs, along with Ryp1 and a second regulator whose motif we report here, guide the expression of endospore-associated transcripts and secreted effectors, including proteases. Ryp1 also controls a core regulon that is expressed in both spherules and hyphae via a canonical Ryp1 motif. Finally, in arthroconidia, Ryp1 impacts the transcriptome, but likely through indirect regulation of one or more additional key regulators.

In particular, S-*RYP1*-dependent genes are more likely to have more Ryp1 motif hits per promoter than any of the other gene subsets we studied, which raises the interesting possibility of cooperativity of Ryp1 binding in spherule conditions. Cooperative binding [83] and stochastic switching [61, 64, 84] have been studied for Wor1, the Ryp1 ortholog in *Candida* species, where Wor1 regulates the white/opaque switch by promoting a specific developmental program. Since Ryp1 both induces and represses the abundance of various transcripts, we hypothesize the existence of additional co-regulators that help mediate the directionality of its regulation in addition to characteristics of the motif’s strength, number, and location within the promoter itself. One of these co-regulators may bind to the second motif discovered to be enriched only in S- *RYP1*-dependent genes, and the motif sequence could be leveraged to discover the identity of that second regulator. The group of candidate transcription factors we have presented here can serve as a roadmap for discovering these additional major co-regulators of the morphologic switch in *Coccidioides*.

### Secreted proteases warrant further study in *Coccidioides*

As we have demonstrated, secreted proteases exhibit intriguing dynamic expression during *Coccidioides* spherulation and comprise an outsized membership of the stringent sets of genes we selected based on our transcriptomic data. Three of the 20 endospore-enriched genes we define are secreted proteases. Additionally, six of the 16 putative secreted effectors we defined are also secreted proteases. Prior genomic sequence analysis has found that two secreted protease families, the S8 serine proteases [57] and the M35 metalloproteases [85], are expanded in *Coccidioides* relative to other fungi, with the M35 family also undergoing positive selection. Given the 9 secreted proteases we report as endospore-associated or putative effectors and a known role for one of these proteases, *MEP1*, in mediating host-endospore interactions [71], it is intriguing to hypothesize that these secreted proteases are involved in *Coccidioides* virulence and could be a large part of its effector armamentarium. Secreted proteases often evoke eosinophilia [86], which is a part of *Coccidioides*’ clinical presentation [87]. It is not yet known whether eosinophils are protective during *Coccidioides* infection. However, given their association with Th2-dominant immune responses that are often not protective in fungal infections [88], an intriguing hypothesis is that the numerous secreted proteases in *Coccidioides* bias toward an ineffective immune response. Interrogation of protease function via mutant generation will be key to elucidating their role in *Coccidioides* virulence.

## Supporting information

Ryp1 deletion mutant genome assembled by pilon

## Acknowledgements

This research was supported by the HHMI Hanna Gray Fellowship (to CH), the Program for Breakthrough Biomedical Research, which is partially funded by the Sandler Foundation (to CH), NIH R21AI172185 (to AS), NIH 5R01AI146584 (to AS) and NIH U19AI166798 (to AS) for funding. AS is a Chan Zuckerberg Biohub – San Francisco Investigator. We acknowledge the UCSF PCAT for use of equipment, UCSF CAT and CZI Biohub San Francisco for sequencing resources, and the Cellular and Molecular Medicine Electron Microscopy Core (UCSD-CMM-EM Core, RRID: SCR_022039) for electron microscopy services. We thank Dr. Sinem Beyhan for generation of the Ryp1 polyclonal antibody used in these studies. The funders had no role in study design, data collection and interpretation, or the decision to submit the work for publication.

## Notes

### Competing Interest Statement

The authors have declared no competing interest.

