## Supplementary Figure Legends for "Transcriptomic atlas of the morphologic development of the fungal pathogen *Coccidioides* reveals key phase-enriched transcripts"

### Supplementary Materials

#### Supplementary Figure Legends

**Figure S1: Early spherulation cultures are synchronous.** A. Micrograph of a spherulation culture demonstrating all 4 possible morphologies quantified in 1B: spherule, spherule releasing endospores which remain associated, free endospores that have disassociated from the spherule that released them, and hyphae. B. Percentage of cells in culture that are rounded (instead of barrel-shaped arthroconidia) 8 hours post placement in spherulation conditions. n $\geq$ 400 cells, quantified by hand for each sample. Baseline round cells in arthroconidia stock are likely barrel-shaped arthroconidia on end. C. Average spherule diameter at day 6 for each replicate (microns), measured manually in Fiji for $\geq$ 50 spherules per condition. Error bars show standard deviation. D. Pearson correlation coefficients, quantitatively shown by color, comparing all samples to each other (3 replicates at each timepoint). E. Number of significantly-differentially regulated transcripts (2-fold change, FDR 5 % using limma) for each comparison of triplicate samples to the previous timepoint.

**Figure S2: Wildtype develops into different morphologies while *ryp1*Δ is hyphal-locked.** A. Micrographs of fixed samples from each flask at the time of RNA harvest for all replicates of spherule growth. Replicate 1 contains the same images as shown in Figure 2A. Subsequent samples for each replicate taken from the same flask over time. B. Average spherule diameter at day 6 for each wildtype replicate in spherule growth (microns), measured manually in Fiji for at least 1000 spherules per condition. Error bars show standard deviation. C. Number of significantly-differential transcripts (2-fold change, FDR 5 % in limma) for each stated comparison in spherule cultures. D. Micrographs of fixed samples from each flask at the time of RNA harvest for all replicates of hyphal growth. Subsequent samples taken from the same flask over time. Black arrowheads indicate branching hyphae. Replicate 1 contains the same images as shown in Figure 2D. E. Number of significantly-differential transcripts (2-fold change, FDR 5 %) for each stated comparison in hyphal cultures.

**Figure S3: Ryp1 induces spherule-related transcripts and represses hyphal-related transcripts.** A. Arthroconidia in Figure 3 were stored for 1-2 days at 4˚C prior to initiating growth of spherules or hyphae, which has been shown to alter the transcriptome [1]. To determine whether storage conditions affected the *ryp1*Δ arthroconidia in a different manner than wildtype arthroconidia, we repeated a limited spherulation and hyphal time-course with arthroconidia that were germinated immediately after harvest. Bar graph showing number of significantly differential transcripts between wildtype and the *ryp1*Δ mutant at each timepoint specified. Transcripts that are induced by *RYP1* (higher in WT than *ryp1*Δ) are in purple and transcripts that are repressed by *RYP1* (higher in *ryp1*Δ than WT) are in green. The number of *RYP1*-dependent transcripts remained highest in arthroconidia compared to early spherule and hyphal timepoints. B. Scatterplot of $\log_2$ of the ratio of wildtype to *ryp1*Δ (counts per million) in arthroconidia from 3A and S3A. C. Scatterplots comparing ratios of $\log_2$(counts per million) for each transcript. Top row: comparing spherule wildtype/*ryp1*Δ to wildtype spherule/wildtype hyphae at each specified corresponding timepoint. Bottom row: comparing hyphal wildtype/*ryp1*Δ expression to wildtype spherule/wildtype hyphae at each specified corresponding timepoint. D. Pearson correlation values from C graphed over time for the top row of C in yellow and the bottom row of C in blue. E. Expression of the *RYP1* transcripts in arthroconidia, all timepoints of spherule development, and all timepoints of hyphal growth, as $\log_2$(counts per million).

**Figure S4: Reproducible Ryp1 binding by ChIP-Seq reveals multiple motifs.** A. Micrographs of fixed samples from each flask at the time of RNA and DNA harvest for all wildtype replicates of spherule and hyphal growth for paired ChIP-seq/RNA-seq experiment. Subsequent samples for each replicate were taken from the same flask over time. B. Micrographs of fixed samples from each flask at the time of RNA and DNA harvest for all replicates of spherule and hyphal growth for the *ryp1*Δ mutant from the same experiment as in A. C. Barplot demonstrating the proportion of morphology-regulated genes (as defined in 3H for Replicate 1, same comparison for RNA-seq generated from samples in A for Replicate 2) whose promoters have Ryp1 binding by ChIP-seq. D. Schematic demonstrating iterative method used to combine Ryp1 binding peaks found by ChIP-Seq, as described in the methods. E. Previously published Ryp1 binding motif in *Histoplasma* [2]. F. Distribution of the number of Ryp1 motif hits per promoter for the motif in 4G for each subset of genes with Ryp1 motif hits as defined in 4H. G. As in F but now describing distribution of the number of Ryp1 motif hits per promoter for the motif in 4I. H. Bar plot of promoter lengths (basepairs) for each gene subset.

**Figure S5: DMEM and RPMI media used to generate spherules.** A. Micrographs of fixed samples from each replicate on day 3 of spherulation in DMEM + 20 % FBS. Spherules were generated from the same arthroconidia stock and grown in the same conditions as spherule samples described in Figure 2. Spherules were also harvested at the same time as microscopy samples for RNA-seq. B. As in A but spherules were generated from 3 days of growth in RPMI + 10 % FBS.

**Figure S6: Three genes in the six-gene cluster influence arthroconidia-associated pigment.** A. Quantification of the number of visible cell wall layers for wildtype and each *DitClusterΔ* mutant. *: p <0.01, **: p <0.001, by unpaired t-test. B. Schematic of the location of the regions deleted in *DitClusterΔ* and *DitClusterSmallΔ* mutants. C. Top: Pictures of tubes holding spore stocks for the indicated genotypes demonstrating their color. Middle: Gaussian blur applied to each picture of the tubes to create a uniform color. Bottom: CMYK color parameters for the center of each Gaussian blur, quantifying the difference in yellow pigment. D. Hyphal radial growth at 30˚C for wildtype and each *DitClusterΔ* mutant. E. Micrographs of fixed samples of spherule development for wildtype and each *DitClusterΔ* mutant. F. As in D for wildtype and each *DitClusterSmallΔ* mutant. G. As in E for wildtype and each *DitClusterSmallΔ* mutant.

**Table S1: Sequences of reagents used for genetic manipulation of *Coccidioides***
Primer sequences are listed in the first 19 rows and were ordered from IDT. Crispr Alt-R crRNA refers to the protospacer sequence used to order CRISPR-Cas9 crRNAs from IDT. Repair cassette sequences indicate the full DNA molecule synthesized by Azenta and used as the template for homologous repair during transformation of *Coccidioides*.

**Table S2: Comma-delimited text file containing transcript abundance over the course of spherulation, related to Figure 1**
Each row corresponds to a transcript. The columns are as follows: UNIQID: systemic gene name from Silveira genome [3]. Systematic gene names with _1, _2 appended have multiple isoforms as detected by kallisto although not all isoforms passed read count filter. NAME: Short gene name from Mandel et al [4]. Cp_anno: GenBank *Coccidioides posadasii* Silveira annotation. CiRS: systematic CiRS gene name for the InParanoid-mapped *C. immitis* RS ortholog. CiRS_anno: Genbank annotation for CiRS ortholog. HcG217B: systematic HcG217B GSC gene name for the InParanoid-mapped *Histoplasma* G217B ortholog. HcG217B_anno: GSC annotation for HcG217B ortholog. The next 13 columns give limma adjusted p-values for differential expression for the listed contrasts. The next 24 columns give kallisto mean row normalized counts for each sample (3 replicates per timepoint of spherulation). The last 13

columns give limma generated $log_2$ fold change values for the listed contrasts. Spores refer to arthroconidia.

**Table S3: Comma-delimited text file containing transcript abundance over the course of spherulation and hyphal growth for wildtype and *ryp1*Δ, related to Figure 2**

Each row corresponds to a transcript. The columns are as follows: UNIQID: systemic gene name from Silveira genome [3]. Systematic gene names with _1, _2 appended have multiple isoforms as detected by kallisto although not all isoforms passed read count filter. NAME: Short gene name from Mandel et al [4]. Cp_anno: GenBank *Coccidioides posadasii* Silveira annotation. CiRS: systematic CiRS gene name for the InParanoid-mapped *C. immitis* RS ortholog. CiRS_anno: Genbank annotation for CiRS ortholog. HcG217B: systematic HcG217B GSC gene name for the InParanoid-mapped *Histoplasma* G217B ortholog. HcG217B_anno: GSC annotation for HcG217B ortholog. The next 57 columns give limma adjusted p-values for differential expression for the listed contrasts. The next 84 columns give kallisto mean row normalized counts for each sample (3 replicates per timepoint of spherulation or hyphal growth for the wildtype and *ryp1*Δ mutant each). The last 57 columns give limma generated $log_2$ fold change values for the listed contrasts. For sample and comparison labels: Spores refer to arthroconidia, “Sil” refers to the Silveira wildtype, Ryp1 refers to the *ryp1*Δ mutant, “spherule”/”myc” indicate spherule and hyphal morphology, and “eighth” refers to the 8 hour timepoint of either spherulation or hyphal growth.

**Table S4: Number of genes with persistent Ryp1 binding in their promoter**

Rows correspond to sample type where WT is wildtype and Ryp1 is the *ryp1*Δ mutant at the listed timepoints in either spherule development (“Spher”), hyphal development (“Hyph”), or arthroconidia (“Arth”). The left side of the table lists the number of peaks found by MACS per each replicate for each sample type. The right side of the table shows the number of genes that had at least 1 Ryp1 binding peak located within their promoters and then a tally of the number of genes whose promoters contained at least 1 Ryp1 binding peak in 2 of the 3 total replicates. Samples highlighted in yellow were those for which we continued downstream analysis.

**Table S5: Comma-delimited text file containing transcript abundance over the course of spherulation and hyphal growth from the same samples used for Ryp1 ChIP-Seq, related to Figure S4**

Each row corresponds to a transcript. The columns are as follows: UNIQID: systemic gene name from Silveira genome [3]. Systematic gene names with _1, _2 appended have multiple isoforms as detected by kallisto although not all isoforms passed read count filter. NAME: Short gene name from Mandel et al [4]. Cp_anno: GenBank *Coccidioides posadasii* Silveira annotation. CiRS: systematic CiRS gene name for the InParanoid-mapped *C. immitis* RS ortholog. CiRS_anno: Genbank annotation for CiRS ortholog. HcG217B: systematic HcG217B GSC gene name for the InParanoid-mapped *Histoplasma* G217B ortholog. HcG217B_anno: GSC annotation for HcG217B ortholog. The next 17 columns give limma adjusted p-values for differential expression for the listed contrasts. The next 36 columns are kallisto mean row normalized counts for each sample (3 replicates per timepoint of spherulation or hyphal growth for the wildtype and *ryp1*Δ mutant each). The last 17 columns give limma generated $\log_2$ fold change values for the listed contrasts. For sample and comparison labels: “Arth” refer to arthroconidia, “WT” refers to the Silveira wildtype, Ryp1 refers to the *ryp1*Δ mutant, “spher”/”hyph” indicate spherule and hyphal morphology.

**Table S6: *Coccidioides* candidate transcription factors**

Each row corresponds to a predicted transcription factor. The columns are as follows: UNIQID: systemic gene name from Silveira genome [3]. NAME: Short gene name from Mandel et al [4]. CiRS: systematic CiRS gene name for the InParanoid-mapped *C. immitis* RS ortholog. HcG217B: systematic HcG217B GSC gene name for the InParanoid-mapped *Histoplasma* G217B ortholog. TF_domains: the Pfam transcription factor domains found in each transcription factor candidate. The next 28 columns are kallisto mean row normalized average counts for three replicates corresponding with each RNA-seq sample (same data as Table S3).

**Table S7: Candidate endospore-related genes in *Coccidioides***

Each row corresponds to a predicted transcription factor. The columns are as follows: UNIQID: systemic gene name from Silveira genome [3]. Systematic gene names with _1, _2 appended have multiple isoforms as detected by kallisto although not all isoforms passed read count filter. NAME: Short gene name from Mandel et al [4]. Cp_anno: GenBank *Coccidioides posadasii* Silveira annotation. CiRS: systematic CiRS gene name for the InParanoid-mapped *C. immitis* RS ortholog. CiRS_anno: Genbank annotation for CiRS ortholog. HcG217B: systematic HcG217B GSC gene name for the InParanoid-mapped *Histoplasma* G217B ortholog. HcG217B_anno: GSC annotation for HcG217B ortholog. The next 69 columns are kallisto mean row normalized counts for each indicated sample. Samples are indicated to be from data presented in Figure 1, 2, or S5. Sample names are defined in Table S2 for data from Figure 1 and Table S3 for data from Figure 2.

**Table S8: Candidate virulence effectors in *Coccidioides***

Each row corresponds to a predicted effector. The columns are as follows: UNIQID: systemic gene name from Silveira genome [3]. Systematic gene names with _1, _2 appended have multiple isoforms as detected by kallisto although not all isoforms passed read count filter. NAME: Short gene name from Mandel et al [4]. Cp_anno: GenBank *Coccidioides posadasii* Silveira annotation. CiRS: systematic CiRS gene name for the InParanoid-mapped *C. immitis* RS ortholog. CiRS_anno: Genbank annotation for CiRS ortholog. HcG217B: systematic HcG217B GSC gene name for the InParanoid-mapped *Histoplasma* G217B ortholog. HcG217B_anno: GSC annotation for HcG217B ortholog. The next 45 columns are kallisto mean row normalized counts for each indicated sample. Sample names are from Table S3.

**Table S9: Ryp1 regulation of dityrosine cluster transcripts**

Each row represents a transcript from the six-gene dityrosine cluster. Each column indicates a timepoint of spherulation/hyphal growth. At each timepoint, expression of the transcript was compared between WT and *ryp1*Δ. “N” means no significant difference between the level of the transcript in WT compared to the *ryp1*Δ mutant at that timepoint. "-" means the transcript level in *ryp1*Δ is significantly higher than in WT. "+" means the transcript level in WT is significantly higher than in the *ryp1*Δ mutant.
