## Supplementary figures and images for "Transcriptomic atlas of the morphologic development of the fungal pathogen *Coccidioides* reveals key phase-enriched transcripts"

### Supplementary Figures

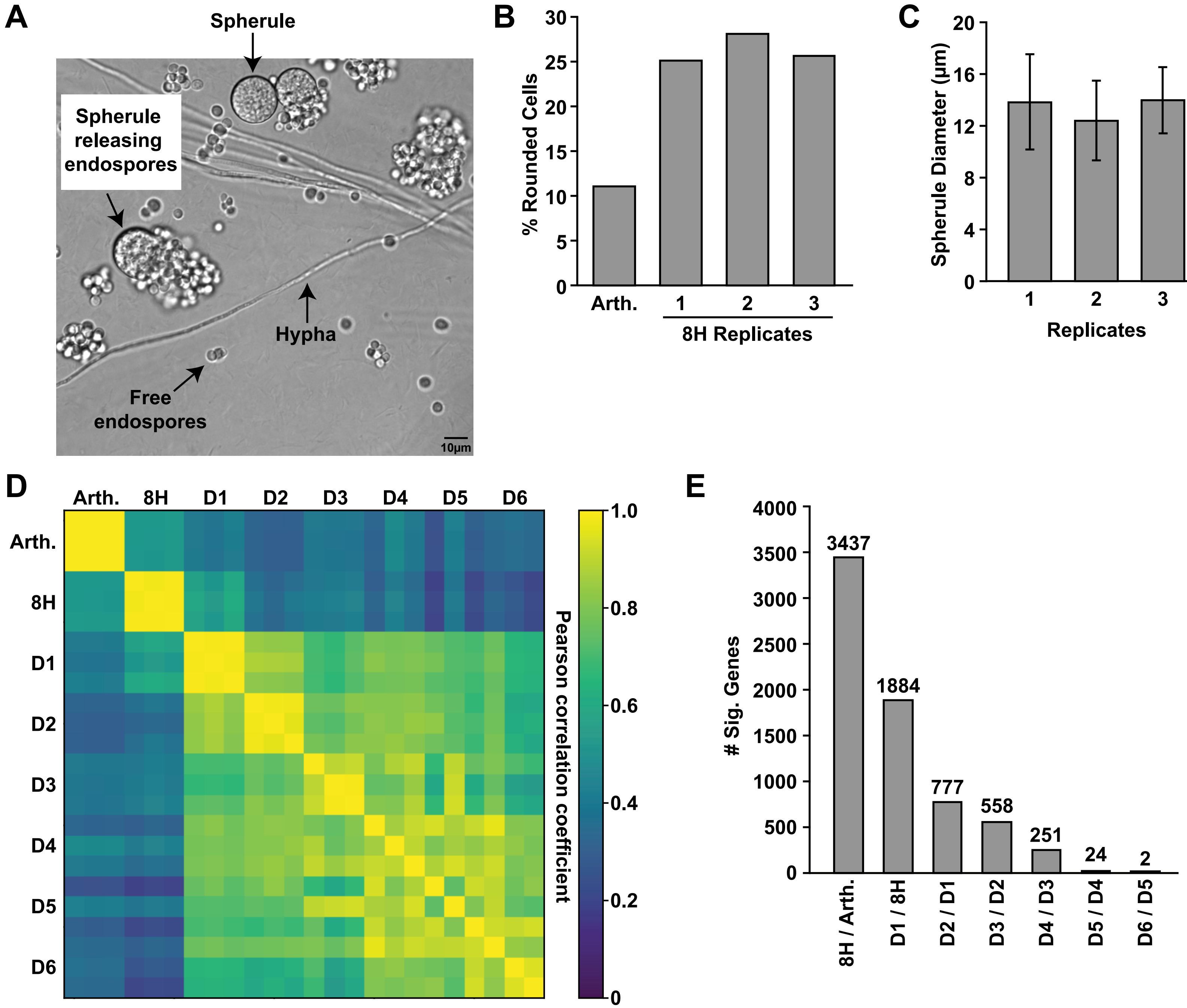

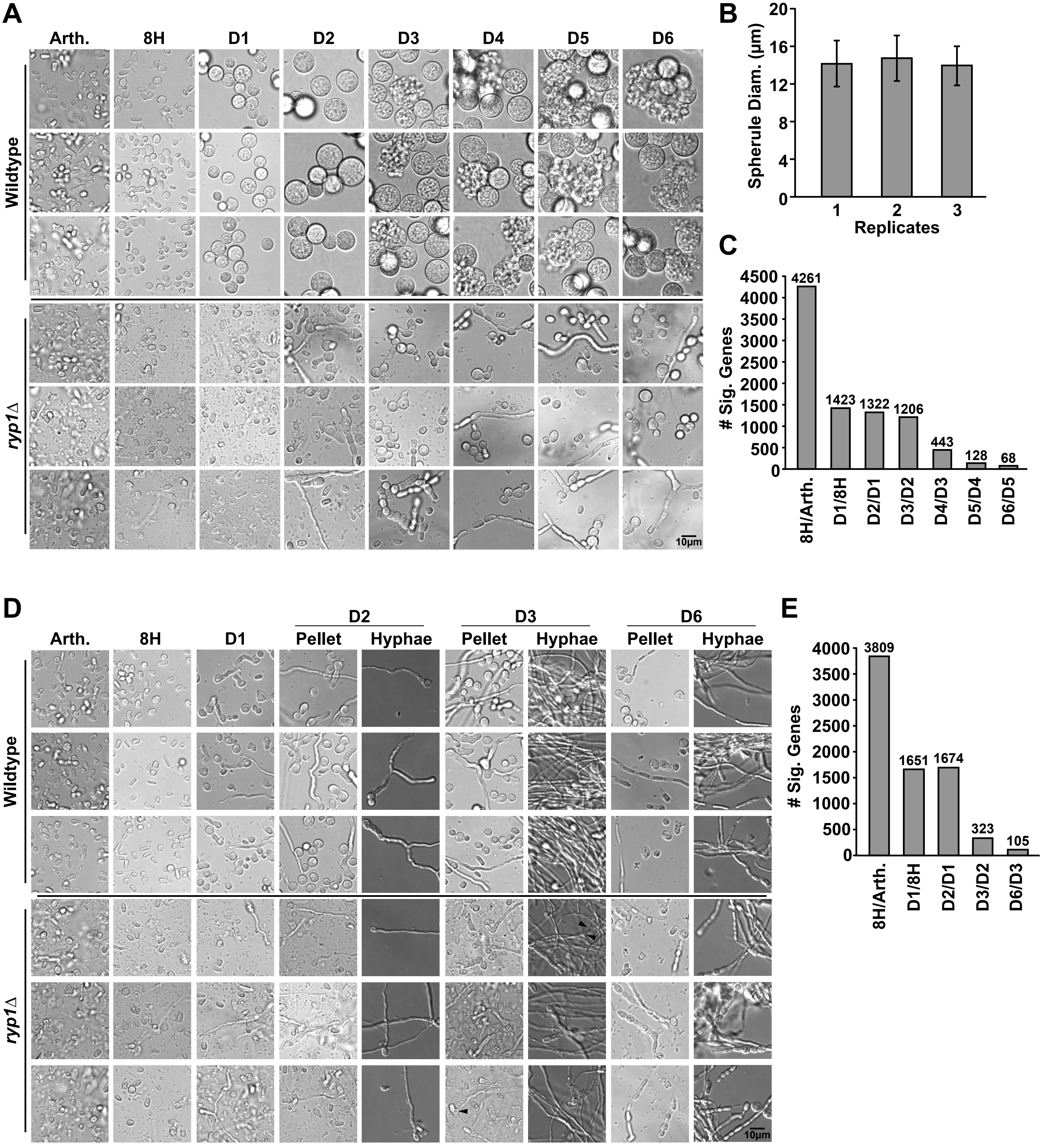

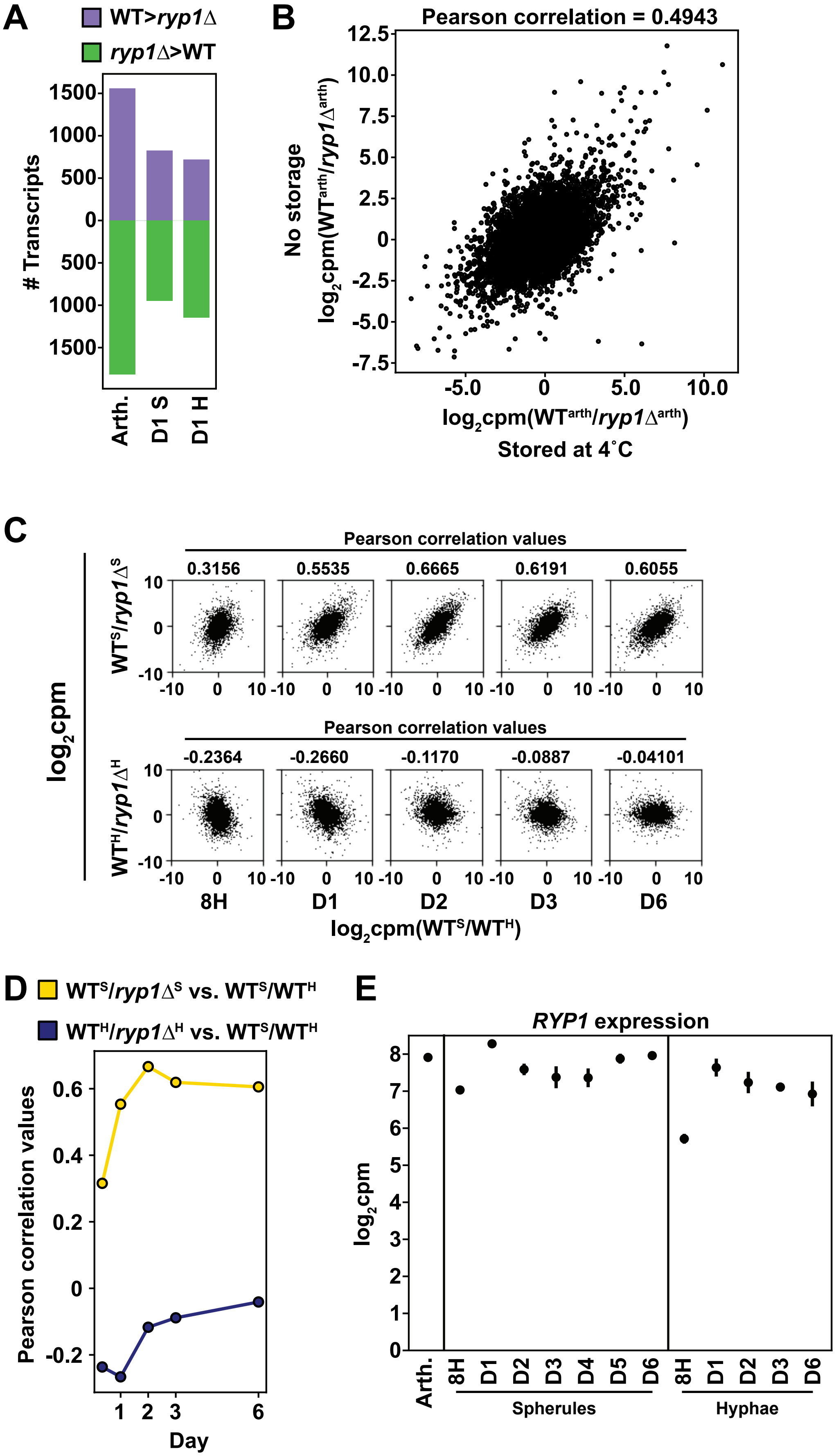

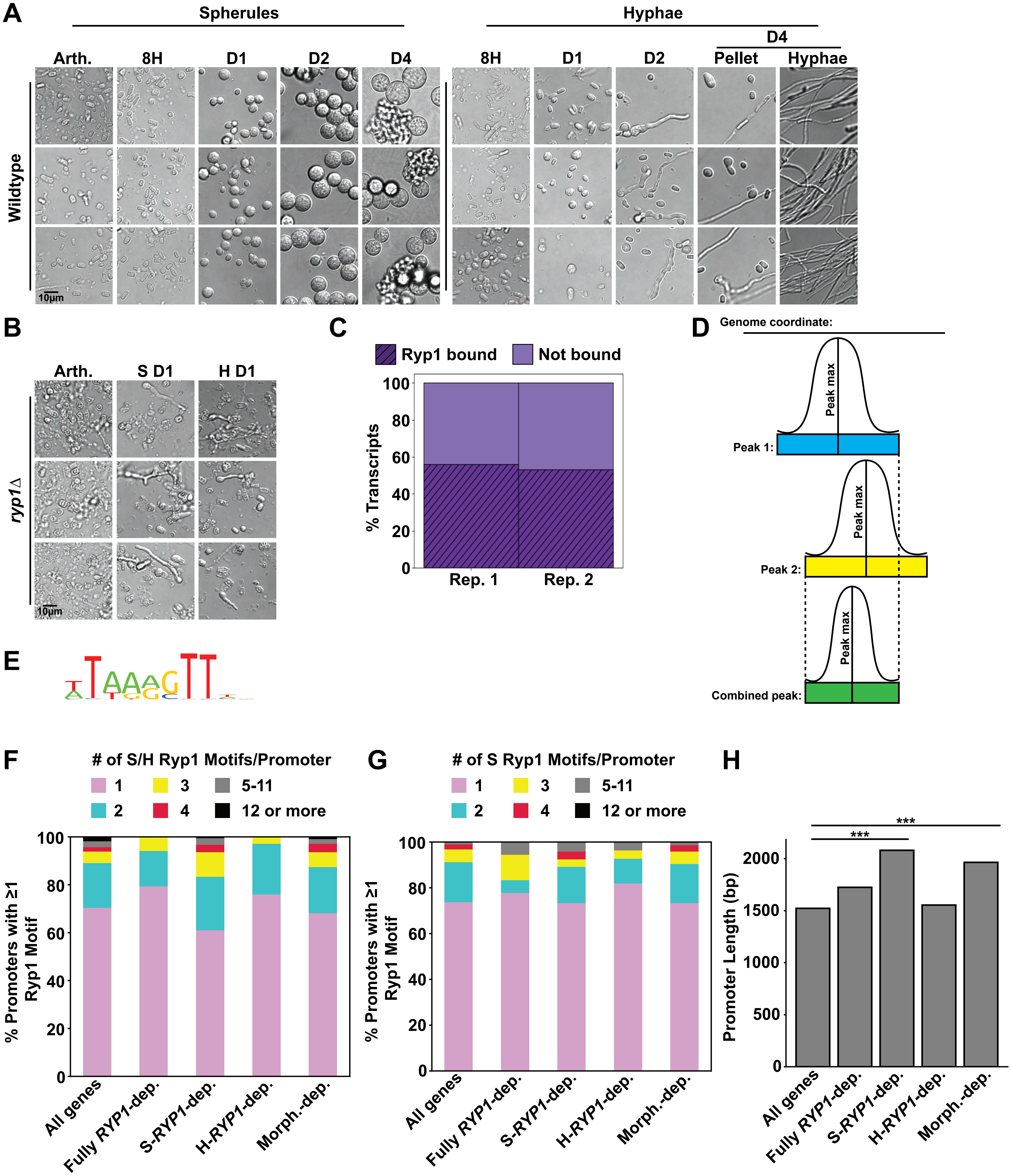

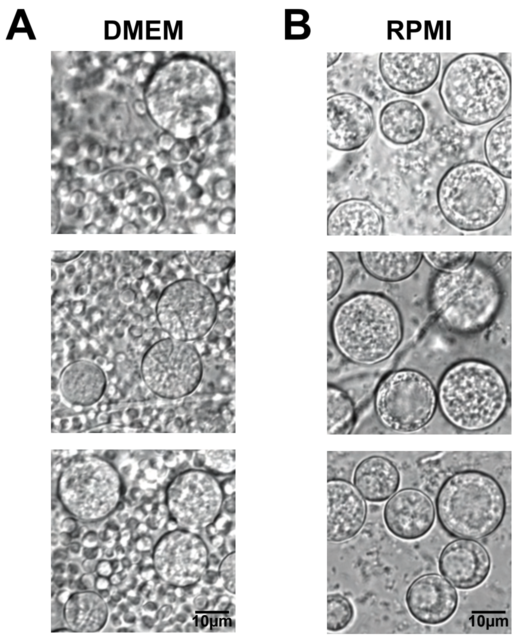

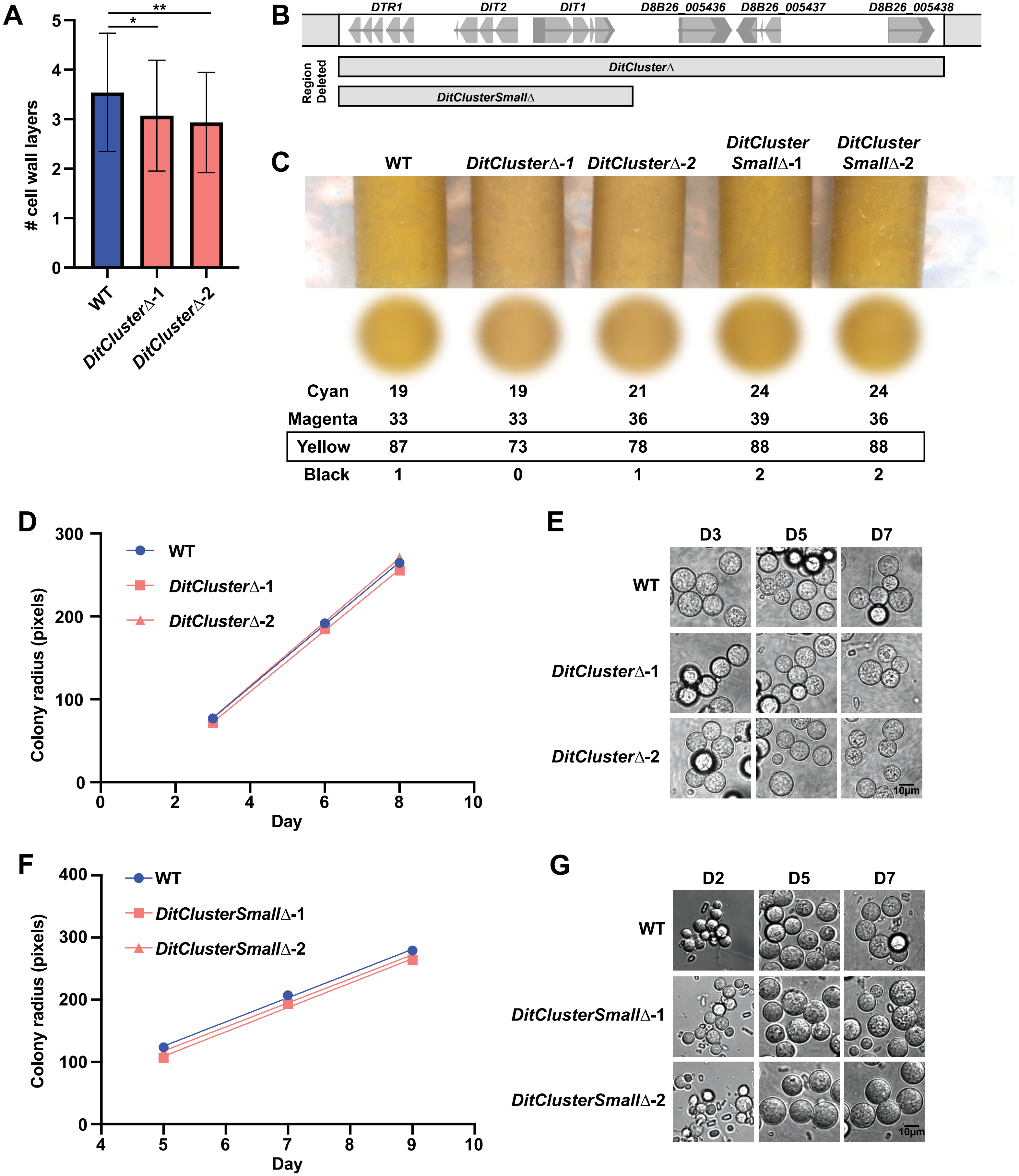
